# 2D Boron Nanoplatelets as a Multifunctional Additive for Osteogenic, Gram-Negative Anti-Microbial and Mechanically Reinforcing Bone Repair Scaffolds

**DOI:** 10.1101/2025.09.05.673512

**Authors:** Jack Maughan, Harneet Kaur, Lucy Prendeville, Tian Carey, Cian O’Connor, Kevin Synnatschke, Juan Carlos Palomeque, Ian Woods, Fergal J. O’Brien, Jonathan N. Coleman

## Abstract

Two-dimensional boron offers unique advantages in bone tissue engineering, unlocking capabilities that conventional additives struggle to achieve. In this study, we leverage the 2D morphology and intrinsic bioactivity of boron nanoplatelets, incorporated into collagen-based scaffolds, to simultaneously achieve osteogenic, neurogenic, angiogenic, anti-inflammatory, mechanically reinforcing, and anti-microbial effects. We synthesize boron nanoplatelets from non-layered precursors using liquid-phase exfoliation and combine them with collagen to form boron- collagen scaffolds (BColl). Boron significantly reinforces the collagen matrix, beneficial for mechanoresponsive bone cells. Osteoblasts and mesenchymal stem cells exhibit healthy morphology and proliferation on BColl films and scaffolds, with extended culture leading to increased alkaline phosphatase release and significantly increased calcium deposition, indicating enhanced osteogenesis. E. coli viability decreases significantly on BColl films, demonstrating their potential to limit post-implantation infections. Finally, we observe angiogenic, neurogenic and anti-inflammatory effects, with dose-dependent upregulation of vascular endothelial growth factor-A, nerve growth factor-beta and interleukin-10, and downregulation of interleukin-6 highlighting boron’s potential to drive pro-reparative processes. Taken together, these data showcase boron’s potential in developing next-generation bone biomaterials, by offering multifunctional benefits to clinically relevant aspects of bone regeneration such as mineralization, angiogenesis, and innervation, while improving the mechanical and anti-microbial properties of natural polymer scaffolds.

**Graphical Abstract & ToC Text:** An ideal bone scaffold would enhance osteogenesis, angiogenesis, and neurogenesis, while preventing inflammation, infection, and stiffness mismatch. 2D materials unlock diverse properties arising from the nanoplatelet morphology, while simultaneously leveraging the intrinsic properties of the material, enabling such multifunctional scaffolds. In this study, we combine 2D boron nanoplatelets with a bioactive collagen matrix to form a multifunctional, versatile bone repair scaffold with osteogenic, angiogenic, neurogenic, anti-inflammatory, and anti-microbial behaviour.

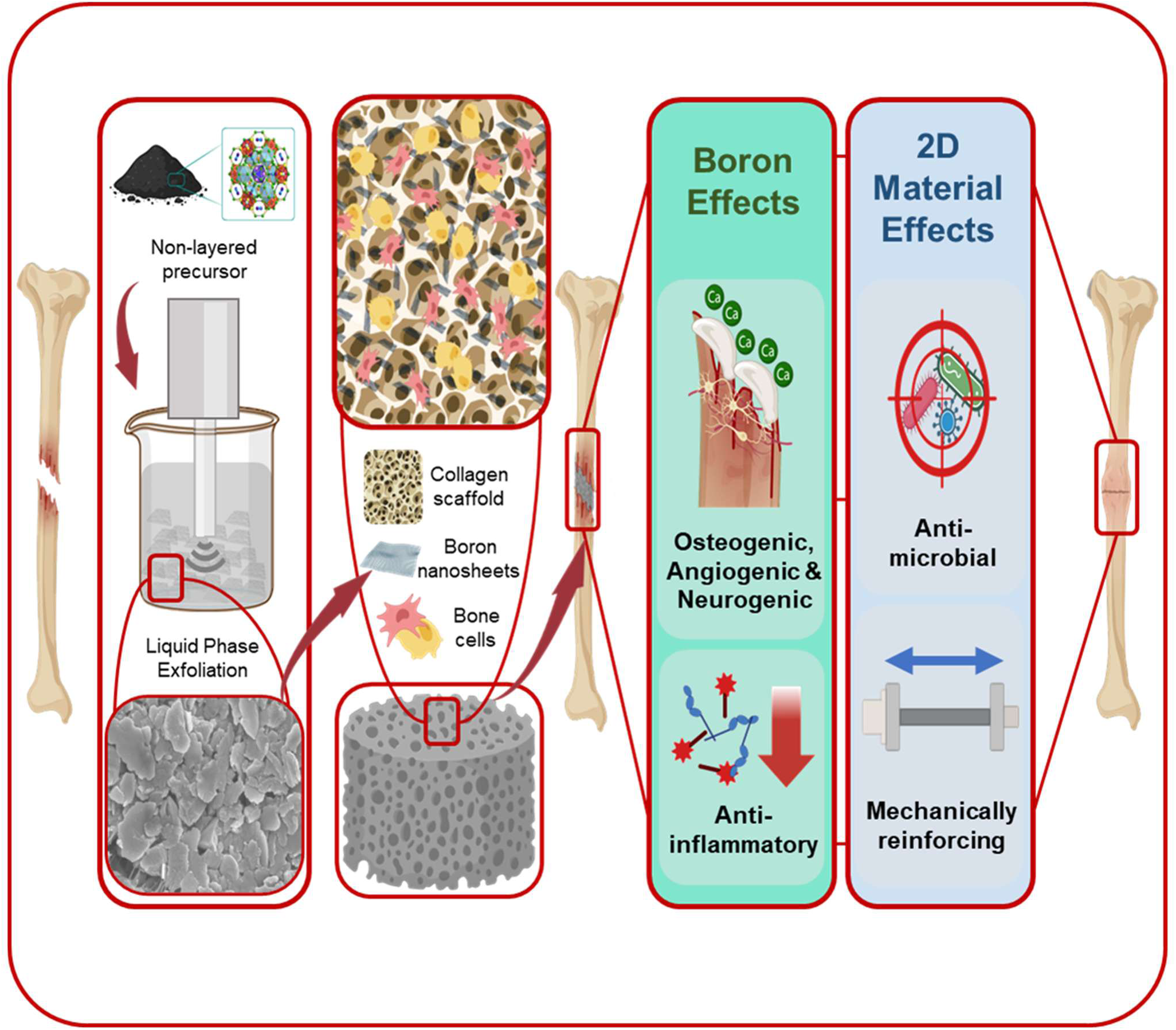

## 1 Introduction

Due to its low cost, scalability, high yield, and versatility, liquid phase exfoliation (LPE) has emerged as one of the primary methods to produce two-dimensional (2D) materials from a wide range of layered materials.^[1]^ The resultant materials have been used in a wide range of applications including electronics^[2]^, batteries^[3]^, coatings & lubricants^[4]^, energy production^[5]^, catalysis^[6]^ and biomedical applications^[7–9]^. Due to the weak van der Waals bonding between layers, it is unsurprising that layered materials such as graphene, MoS2 and h-BN can be exfoliated in this way.^[10]^ What is much more surprising is that LPE can be used to exfoliate non-layered materials, which contain strong chemical bonds in all three dimensions.^[11]^ In such materials, the limited spatial anisotropy in the chemical bonding makes exfoliation considerably more challenging, and would be expected to result in relatively low aspect ratio nanoparticles.^[11]^ These issues can be countered to a certain extent by careful selection of the material to be exfoliated, as certain crystals exhibit some anisotropy in the bonding strength of their lattice planes. The resultant preferential cleavage can lead to the production of quasi-2D nanoplatelets with somewhat enhanced aspect ratios. Various papers have reported nanoplatelets produced in this way, with aspect ratios that can range from as low as 4 to as high as 1400.^[11,12]^ The exact mechanism of this surprising result is highly material dependent, can be tuned using techniques such as cryogenically-mediated exfoliation,^[13]^ and appears to be driven by the formation of cracks at intrinsic defects in a bulk precursor,^[12]^ which then propagate through the material. Judicious choice of solvent and surfactant or polymer stabilizers is also essential for preventing re-aggregation of exfoliated materials, while enhancing hydrophilicity and, if desired, biocompatibility.^[14]^

Developments in the field of exfoliation of non-layered materials have dramatically increased the number of materials that could be exfoliated and subsequently exploited in their 2D form. This opens up a number of previously inaccessible application areas, including a wide range of biomedical applications. For example, boron is a key non-layered trace element in the human body, positively influencing bone health and osteogenesis,^[15–20]^ angiogenesis^[21,22]^, and wound healing,^[23]^ among many other biological systems.^[24,25]^ Indeed, while boron nanoparticles have been synthesized and used in a variety of applications,^[26]^ research into other dimensionalities of nanoscale elemental boron has only picked up speed in recent years, as production of 2D nanoplatelets from non-layered boron is not trivial. While substrate-supported borophenes of the β12, χ3 and 8-Pmmn allotropes, and even some in freestanding form,^[27–37]^ have been synthesized using chemical vapor deposition, molecular beam epitaxy and mechanical exfoliation, these are expensive, time consuming, and low yield synthesis techniques.^[29,35,38]^ Recently, however, top-down approaches such as LPE have demonstrated efficient exfoliation of boron nanoplatelets consisting of β-rhombohedral boron, with the same crystallinity as the bulk precursor.^[12,36,39–47]^ Systemic boron supplementation has been studied for various beneficial health outcomes,^[18,24]^ along with boron-containing compounds successfully being used for bone tissue engineering applications, leading to enhanced bone repair and mineralization.^[17,48–50]^ Boron nanosheets, however, have yet to be applied to the repair of large bone defects - this is a challenging prospect, since robust osteogenesis, angiogenesis and innervation are required to direct sustainable bone repair.^[51]^ On top of this, the risk of infection,^[52]^ the inflammatory response of the tissue,^[53]^ and the stiffness of the biomaterial^[54]^ must be suitably managed to direct cell responses and prevent complications. Despite its difficulty, the repair of large bone defects is highly sought after, as current approaches to bone tissue regeneration struggle with defects above a critical size, representing a significant unmet clinical need.^[55]^

2D materials are particularly promising for tissue engineering thanks to unique properties such as high surface area-volume ratio,^[56]^ sharp edges,^[57]^ and network effects on the mechanical properties of composites.^[58]^ These properties, which arise directly from their 2D nature and are not found in their bulk counterparts, unlock a diverse range of possibilities and applications, from anti-microbial effects^[59]^ and drug delivery,^[60]^ to mechanical reinforcement.^[58]^ Building on previous experience in both liquid phase exfoliation of non-layered nanomaterials,^[11,61]^ and in the development of 2D nanoparticle composites,^[62,63]^ we aimed to incorporate 2D boron within a macroporous, biocompatible and biodegradable collagen scaffold^[64,65]^ and test its multifunctional pro-regenerative characteristics for bone tissue engineering applications. Since natural polymers excel biologically as scaffold materials, but often display weaker mechanical properties,^[64,66]^ additives such as hydroxyapatite^[67,68]^ and carbon nanomaterials^[67,69]^ are often used to strengthen the material, taking advantage of the preference of mechanosensitive bone cells for stiffer substrates, and the importance of mechanical properties in many bone repair applications.^[70]^ To reduce the major problem of bone infection,^[71]^ metal ions, antibiotics, and chitosan are sometimes used to impart anti-microbial activity.^[52,72,73]^ To enhance the regenerative capacity of the materials, further drug and gene delivery is often carried out using nanoparticle carriers,^[60,74]^ while osteogenic, angiogenic, and neurogenic effects are often enhanced by various bioactive ions, materials or growth factors such as Mg^2+^, hydroxyapatite or the addition of recombinant growth factors such as BMP-2.^[75–79]^ Each of these interventions can enhance the potential of natural polymer scaffolds for bone tissue repair. However, few materials have demonstrated the multifaceted capabilities of 2D boron, which we propose can potentially leverage both the intrinsic properties of elemental boron for osteogenesis, angiogenesis, and neurogenesis, and the emergent effects of 2D materials for mechanical reinforcement and anti-microbial activity. We therefore propose that 2D boron is an ideal multi-functional additive for bone tissue engineering.

To demonstrate this potential, the specific aims of this study were to first synthesize boron nanosheets from a bulk precursor by LPE, followed by their physical and biological characterisation. The study next aimed to fabricate boron-collagen (BColl) composite films and 3D scaffolds by lyophilization, characterize their mechanical properties, and assess the osteogenic potential of these 3D scaffolds with physiologically relevant bone cells. Finally, the study endeavored to demonstrate the functional effects of this material by assessing the anti-microbial, angiogenic, neurogenic, anti-inflammatory, and mechanically reinforcing effects of boron loading.

## 2 Results and Discussion

### 2.1 Synthesis of boron nanoplatelets

Boron is a relatively new member of the two-dimensional material family due to its non-layered nature.^[80]^ As a result, before commencing biological characterisation, optimization of the synthesis protocol was required, in order to both achieve optimal flake yield and quality, and high biocompatibility by means of non-cytotoxic stabilizing molecules. This optimization was carried out along three axes – bulk crystallinity, solvent choice, and exfoliation method. With amorphous boron powder as the precursor, liquid phase exfoliation was carried out in a range of solvents - isopropanol (IPA), ethanol (EtOH) and N-methyl-2-pyrrolidone (NMP). These initial exfoliations failed to yield particles with a 2D morphology, instead yielding 0D nanoparticle dispersions, as seen in **Figure S1a-c**. This result suggests that the lack of anisotropy in boron bonding leads to isotropic cleavage, and the subsequent formation of nanoparticles. However, the efficiency of LPE for non-layered materials can be enhanced in similar ways to that of LPE for layered materials. For example, cryogenic treatment can be used to render inter-atomic bonds more brittle, and thus prone to fracture during exfoliation.^[11,81]^ In order to induce this thermal fracturing, and thus exfoliation along a plane, cryogenic exfoliation in IPA and NMP was then tested. This approach was more successful, yielding a mix of morphologies, including 0D nanoparticles, 1D nanoribbons, and 2D nanoplatelets (Figure S1d). These morphologies potentially arise due to the formation of cracks following cryogenic treatment, which then propagate through the material.^[11]^ However, it was not possible to efficiently separate these morphologies, nor to render the exfoliation more selective for 2D nanoplatelets. Based on these results, we shifted to the use of crystalline boron precursors, which provide crystalline planes along which exfoliation can occur,^[11]^ due to intrinsic stacking faults of boron icosahedra in the 3D crystal reducing the cleavage energy along the {001} plane.^[12]^ With this in mind, 2D boron nanoplatelets were ultimately exfoliated successfully from a crystalline precursor in NMP, chosen for its favorable surface tension and solubility parameters.^[1,43]^ As seen in the scanning electron microscopy (SEM) (**Figure 1**a) and transmission electron microscopy (TEM) (Figure 1b) data, the resulting boron nanoplatelets exhibited a 2D morphology indicative of successful selective exfoliation, which was confirmed by atomic force microscopy (AFM) measurements (Figure 1c).

**Figure 1.**
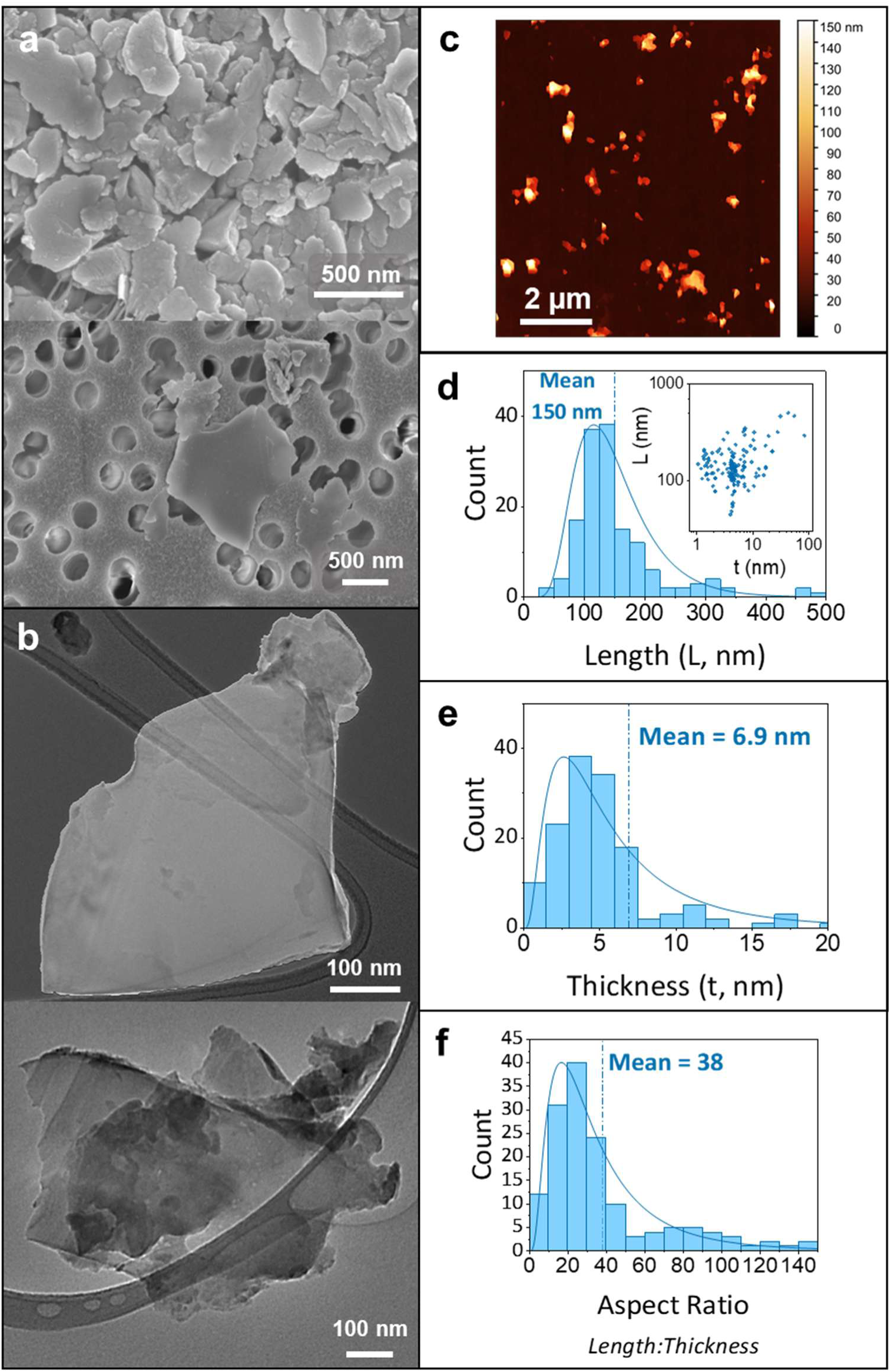
Morphological characterisation of boron nanosheets: Scanning electron microscopy (SEM) (**a**, scalebars 500 nm) and transmission electron microscopy (TEM) (**b**, scalebars 100 nm) imaging were used to visualize successful exfoliation of the boron nanosheets. **c-f)** Atomic force microscopy (AFM) imaging (a) of 147 boron nanosheets was used to assess the length (d), thickness (e) and aspect ratio (f) distributions of the nanosheets, corroborating the 2D morphology seen in the SEM and TEM. Scalebar 2 µm

Statistical analysis of these images revealed a broad distribution of nanoplatelet dimensions, with lengths (L) ranging from 50 to 500 nm and thicknesses (t) ranging from 1 to 80 nm (Figure 1d, inset). The nanoplatelet length and thickness distributions are shown in Figure 1d and e as histograms. The average flake length was measured as 150 ± 6 nm, with an average thickness of 6.9 ± 0.8 nm. Knowledge of nanoplatelet length and thickness allows the calculation of the aspect ratio 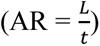. The AR distribution is shown in Figure 1f and has a mean of 38. This data gives us an insight to the surface area:volume ratio of our material, which is critical for many applications^[82]^ - for the average nanoplatelet dimensions attained in this work, the surface area:volume ratio is 0.32, >5 times larger than that of the smallest widely commercially available nanoparticles, which have a radius 50 nm^[83,84]^ (**Figure S5**). This means more of the material is exposed to the surrounding tissue at any given time, improving the efficiency of the material at delivering intrinsic biological effects.^[85]^

### 2.2 Physical characterisation of boron nanoplatelets

Following the successful exfoliation of boron nanoplatelets, the crystal, thermal and optical properties of the material were established. A primary concern about elemental boron materials is oxidation,^[86]^ since it can lead to changes in material properties such as conductivity, surface chemistry and biocompatibility.^[87–89]^ To assess the degree of oxidation suffered by the material, thermogravimetric analysis (TGA) and energy-dispersive X-ray spectroscopy (EDX) were carried out. The tendency of the material to oxidize was first observed in TGA data (**Figure 2**a), which showed that upon heating of boron under an air atmosphere, the mass increased significantly due to the formation of oxides, with the rate of oxidation increasing rapidly beyond 500 °C. The main oxidation peak (Figure 2a, inset) is centered at 550 °C, consistent with previous reports of the oxidization of boron.^[86,90]^

**Figure 2.**
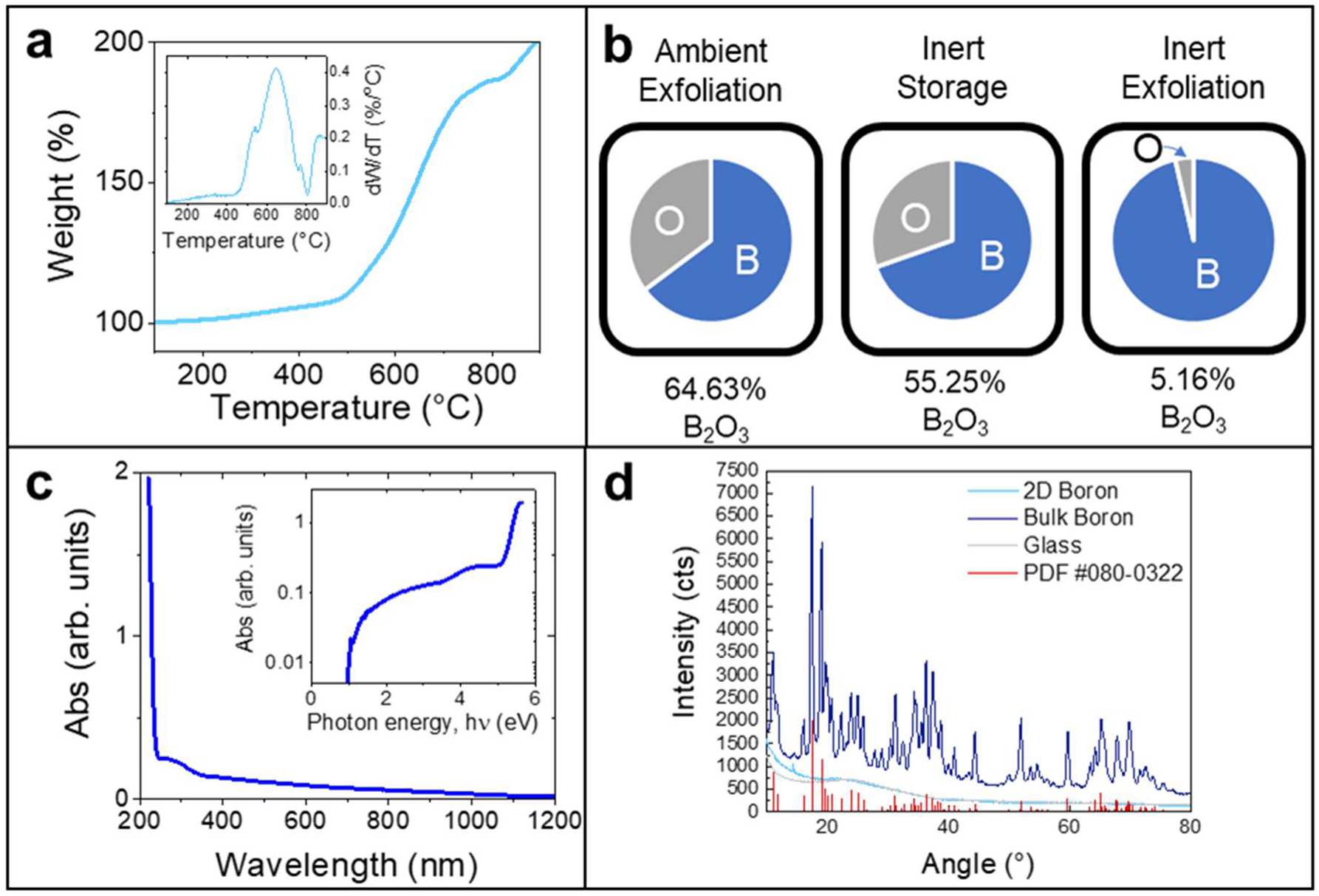
Physical characterisation of boron nanoplatelets: **a)** Thermogravimetric analysis (TGA), with an increase in sample mass with temperature showing oxidation of boron nanoplatelets upon heating. Inset shows derivative of mass loss. **b)** Energy-dispersive X-ray spectroscopy (EDX), used to assess the elemental makeup (atomic %) of the sample under different conditions of exfoliation and storage – inert exfoliation was demonstrated to have the largest impact on prevention of oxidation in the material. **c)** UV Visible light spectroscopy (UV Vis) of a boron nanoplatelet dispersion showing absorption spectra typical of a semiconducting nanomaterial. **d)** X-ray diffraction patterns (XRD) used to assess the crystallinity of the starting bulk boron powder and exfoliated boron nanoplatelets, showing a potential shift towards an amorphous allotrope following exfoliation

EDX (Figure 2b) was then carried out both on samples exfoliated in ambient conditions with commercial solvents, and on samples exfoliated in inert conditions with anhydrous, deoxygenated solvents. Boron exfoliated in both conditions exhibited incomplete oxidation after prolonged storage in ambient and inert conditions, as indicated by comparison with the most common boron oxide, B2O3, which would correspond to 60 at% oxygen.^[91]^ This suggests the occurrence of surface oxidation, slowing oxidation of the bulk of the nanoplatelet. Furthermore, these analyses show that the oxidation of the boron primarily occurs during the exfoliation process as a result of dissolved oxygen in the solvents, rather than during storage, as ambient-exfoliated boron that was stored under inert conditions still exhibited a significant amount of oxidation. As a result of these measurements, the standard exfoliation protocol was adapted to use inert, degassed, and anhydrous solvents. While accurate quantification of oxygen content is challenging due to the limitations of EDX, the relative oxidation of these materials indicates the formation of a stable passivation layer of B2O3 on the surface of the boron nanoplatelets. Furthermore, boron oxide has also been demonstrated to have similar bioactive effects to other forms of boron and boron-containing compounds, meaning it maintains its utility for bone tissue engineering scaffolds.^[92–94]^

The final characterisation of the boron flakes was intended to reveal any shifts in crystallinity or optical properties arising after exfoliation. The UV-visible absorption spectrum of a dispersion of boron nanoplatelets exfoliated in NMP under inert conditions is shown in Figure 2c. We note that the considerable scattering background, typical of nanomaterial dispersions,^[95]^ has been removed (**Figure S3**). The same spectrum has been re-plotted in the inset, with the absorbance on a log scale, plotted versus photon energy. This graph clearly shows our boron nanoplatelets to be semiconducting with a bandgap of slightly under 1 eV. This could be consistent with either crystalline boron or amorphous boron which display optical gaps of 0.9 and 0.7 eV respectively.^[96]^ However, our relatively featureless spectrum is more in line with amorphous boron. The increase in absorbance below 250 nm (above 5.1 eV) may arise due to the presence of boron oxide, an insulator with a larger bandgap, around 5.23 eV.^[97]^

In order to assess whether there was a shift in the crystallinity of the material due to the exfoliation, powder X-ray diffraction (XRD) (Figure 2d) was carried out. The XRD spectrum for the bulk boron precursor is consistent with the crystalline form of β-boron, as shown in the literature,^[43,98]^ and matches well with the relevant powder diffraction file from the ICDD database^[99]^ (PDF #080-032).

The XRD spectrum for the 2D boron nanoplatelets, on the other hand, is indicative of a shift towards an amorphous allotrope of boron/boron oxide. Such amorphization has been observed previously for LPE of non-layered materials such as red phosphorus.^[3]^ Furthermore, this indicates that no transition to a borophene-like phase occurs. This is consistent with previous work on LPE of crystalline boron precursors,^[39,43]^ suggesting that top-down exfoliation of boron leads to the formation of few-layer nanoplatelets consisting of the bulk material, rather than the allotropically distinct borophene phases.^[100]^ Taken together, the physical characterisation of the material shows that we have successfully exfoliated 2D nanoplatelets of amorphous boron. We believe these are suitable for leveraging the benefits of both the 2D morphology and the intrinsic properties of boron.

### 2.3 Fabrication and characterisation of 2D BColl films and 3D BColl scaffolds

We propose that one important application of our boron nanoplatelets is as a filler material in polymer-based composites. One significant advantage of high aspect ratio nanoplatelets, when compared to quasi-spherical nanoparticles, is their ability to mechanically reinforce polymers^[101]^. Stiffness is well-established as a key regulator of cellular differentiation,^[102,103]^ and thus mesenchymal stem cells (MSCs) will exhibit different differentiation behaviour on a stiff substrate than a soft substrate. Models such as shear-lag theory^[104]^ and Halpin-Tsai theory^[105]^ clearly show that composite reinforcement depends sensitively on filler particle aspect ratio, with higher AR particles leading to greater reinforcement.

To test the magnitude of mechanical reinforcement and its effect, composite films were fabricated by blending boron nanoplatelets with collagen to form boron-collagen (BColl) slurry, followed by casting into homogenous films of 0, 0.5, 1, 2.5 and 5 vol% boron loading (**Figure 3**a). Tensile testing was then carried out using thin strips of these BColl films. Examples of stress strain curves for BColl composites with differing boron volume fractions (Vf) show a clear increase in stress at all strains as the boron loading is increased, a clear sign of reinforcement (Figure 3b). While various parameters can be extracted from these stress-strain curves, we focus on the Young’s modulus (E, Figure 3c) and the ultimate tensile strength (UTS, Figure 3d). In both cases, we see steady increases as the volume fraction is increased, with saturation occurring at a loading of roughly 2.5 vol%, and both parameters undergoing a near doubling in their values on addition of boron. The average Young’s modulus of BColl composites was 973 MPa, comparing favorably to the typical stiffness range of trabecular bone, 10 MPa – 3000 MPa.^[106]^ A significant increase was further observed in the initial tensile modulus of the material (**Figure S9a**), within the first 0.5% strain, the regime primarily experienced by cells, as well as a trend towards an increase in the toughness (Figure S9b). These increases render the environment more suitable for osteogenic differentiation, potentially contributing to the increase in osteogenic potential observed for the BColl material.

**Figure 3.**
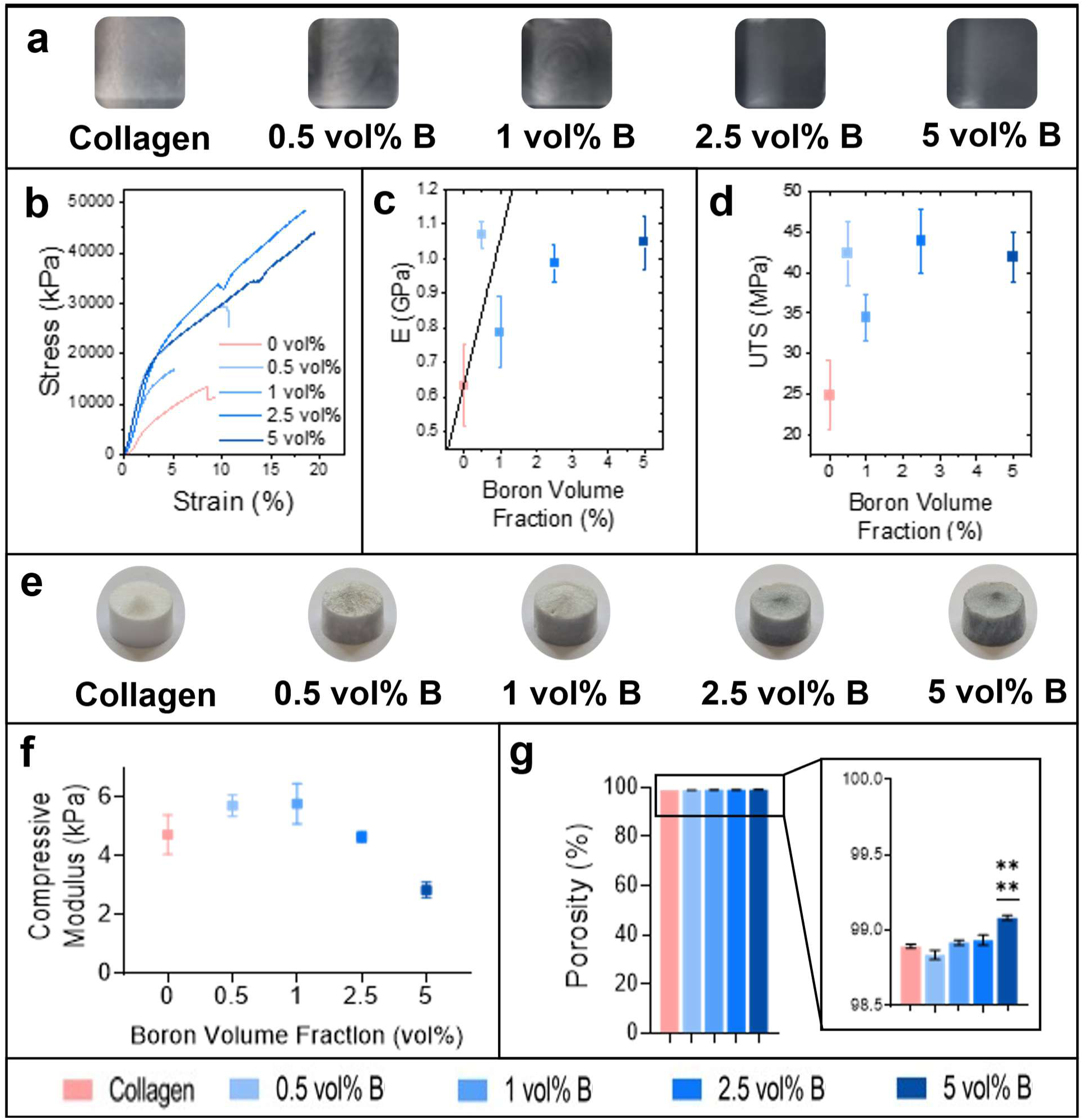
Characterisation of BColl composite films & scaffolds: **a)** Images of BColl films at each boron loading. **b)** Representative stress-strain curves for BColl composites. **c-d)** Measurement of the Young’s modulus (c) and ultimate tensile strength (d) for BColl films, showing significant reinforcement with boron loading. **e)** Images of BColl scaffolds at each boron loading. **f)** Compressive modulus testing of BColl scaffolds. **g)** Measurement of the porosity of BColl scaffolds at varying boron loadings, showing a slight increase in porosity for 5 vol% boron loading. Significances: *p < 0.05, **p < 0.01, ***p < 0.001, ****p < 0.0001.

To understand the reinforcement of the BColl composite films, we analyze the data in Figure 3c using Halpin-Tsai theory.^[105]^ The Halpin-Tsai equations describe the composite modulus, E, as a function of the filler volume fraction, Vf, the moduli of filler, EF, and matrix, EM, as well as the platelet aspect ratio, L/t. This particular model is appropriate when nanosheets are aligned in the plane of the composite, and has been shown to describe composites of BN nanosheets in polymer matrices quite well.^[107]^ <u>_ENREF_12</u>It has been shown that applying the approximations that *E_F_* / *E_P_* >> 1 and*V_f_* << 1, which are appropriate for most nanocomposites, to the basic Halpin-Tsai equation gives the simplified expression^[107]^:

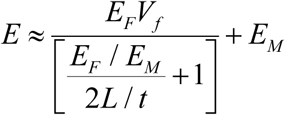

This equation predicts a linear increase in composite modulus with nanoplatelet volume fraction. We can test the applicability of this equation by plotting its prediction on top of our data using our measured values for EM (0.64 GPa) and 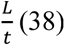, as well as a previously reported value for the stiffness of amorphous boron (EF = 300 GPa).^[108]^ This prediction is shown as the solid line in Figure 3c and matches the data reasonably well, especially at low loading values where the model is most applicable. The data deviates from the model at higher loadings, where Halpin-Tsai theory predicts nanocomposite behaviour less accurately.^[109]^ This clearly shows that our stiffness results are consistent with theory, confirming that boron nanoplatelets are an appropriate reinforcing material for collagen. These results imply that boron nanoplatelets are a promising multifunctional composite component for reinforcing the struts of collagen-based tissue engineering scaffolds. Given the small size of the boron nanoplatelets compared to the collagen microstructure in scaffolds, the tensile modulus observed in Figure 3c can be interpreted as an estimate for the modulus of the collagen struts and films upon and between which cells sit in the scaffold.

To extend this analysis into 3D, macroporous BColl composite scaffolds (Figure 3e) were produced by lyophilization, at boron loadings of 0, 0.5, 1, 2.5 and 5 vol%. Compressive modulus testing of these BColl scaffolds revealed a slight trend towards reinforcement at lower boron loadings (Figure 3f), followed by a drop-off in modulus, likely to be related to the increased porosity of the higher loading scaffolds (Figure 3g) – this increase was observed for the highest boron loading, and it is posited that it is due to a slight dilution of the collagen slurry caused by the addition of the boron dispersion during synthesis.

### 2.4 Biocompatibility of boron/collagen (BColl) composites with bone cells in 2D films

As with any novel biomaterial additive, it was necessary to establish the biocompatibility of the boron nanoplatelets before proceeding with functional testing, to ensure safe degradation and excretion of the scaffold after implantation. Boron nanoplatelets, stabilized with polyvinylpyrrolidone (PVP), a biosafe polymer^[110]^ for biocompatibility and hydrophilicity, were added to the media of osteoblasts (MC3T3-E1, osteogenic cell line^[111]^). Cells were grown in suspension with 40 µg mL^−1^ and 80 µg mL^−1^ of boron for 72 h. Metabolic activity and DNA content analysis (**Figure 4**a) showed robust proliferation of the cells in the presence of boron, with a significant increase in both measures for 80 µg mL^−1^ of boron after 72 h. This indicates that the material is non-cytotoxic and can promote proliferation of bone cells, even in free suspension with high concentrations of the material. This is essential, as during the degradation of BColl scaffolds, boron will be released into the surroundings and must be safely excreted. Once the biocompatibility of the boron nanoplatelets had been established, to further demonstrate that BColl composite materials were well-tolerated by cells, osteoblasts were grown on the surface of 0 vol%, 0.5 vol%, and 1 vol% BColl films for 7 days, similar to previous work in our group assessing the biocompatibility of nanocomposite materials.^[62]^ Metabolic activity and DNA content analysis demonstrated robust proliferation of the cells across the surface of the material (Figure 4b). The cell coverage observed for the BColl materials was significantly higher than that of collagen (Figure 4c), as verified by fluorescent staining of the cells with phalloidin and DAPI, with confluent cell layers forming on the surface of the BColl samples (Figure 4d), showing that BColl is not only well-tolerated in a 2D culture environment with bone cells, but that it appears to be enhancing the proliferation and viability of these cells, due to the bioactive effects of boron and its oxides. Various mechanisms that explain ion-mediated enhancement of growth fail to account for boron, which typically does not exist in the B^3+^ form due to the high ionization energies needed to achieve this state,^[112]^ so other research suggests that the release of bio-active ions from the covalent compounds of boron^[113]^ leads to activation of the BMP and MAPK pathways, and subsequent osteogenic and angiogenic effects such as increased RUNX2 phosphorylation and VEGF upregulation.^[15,114–116]^ In addition to enhancing osteoblast differentiation, boron has been shown to inhibit osteoclastogenesis via the downregulation of osteoclastogenic genes such as RANK and MMP-9 and inhibition of the RANKL/RANK pathway.^[116]^ Finally, boron has been shown to influence macrophage polarization and inhibit the TLR pathway, the immunomodulatory effect of which is beneficial for bone implant materials.^[116]^

**Figure 4.**
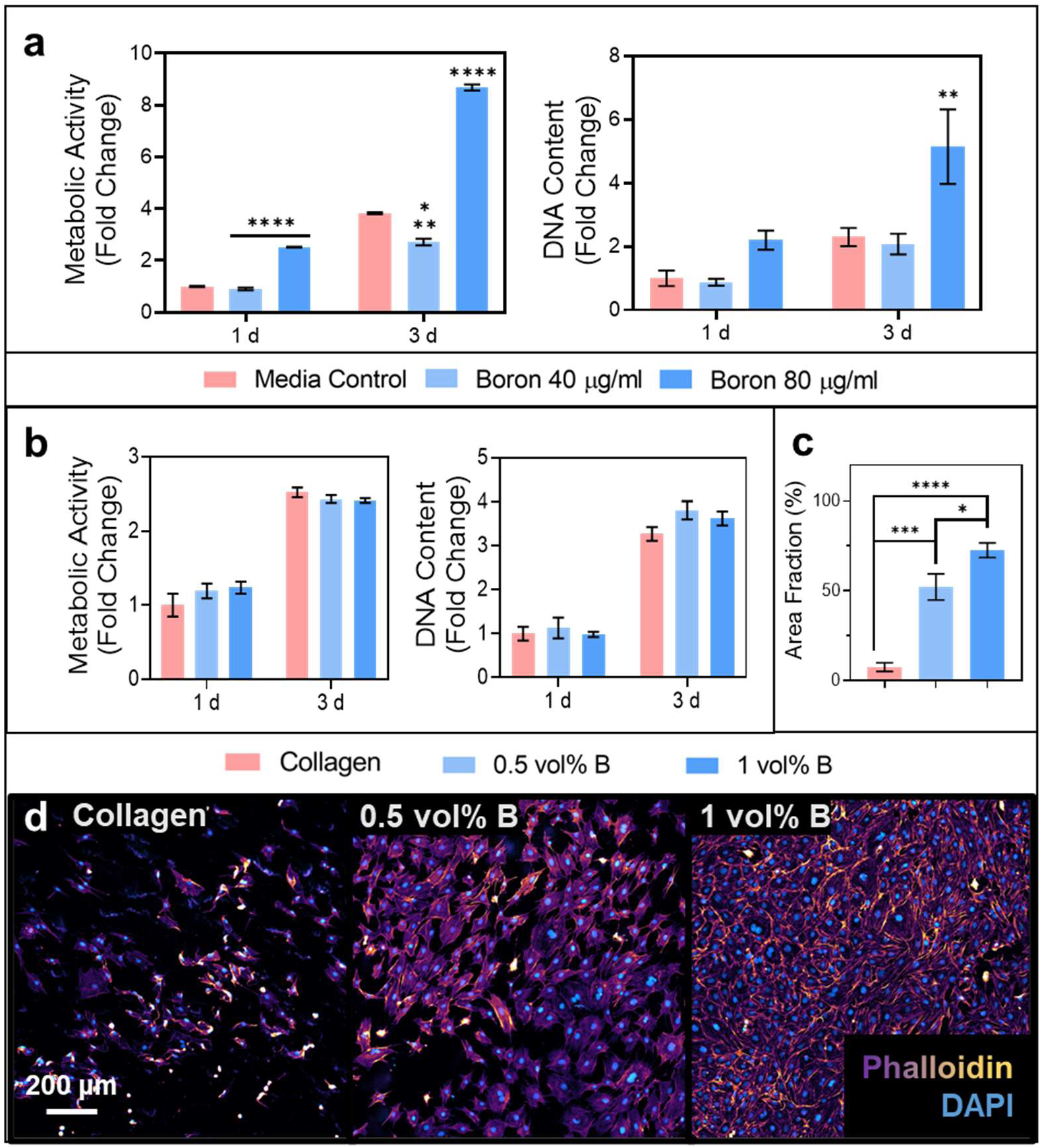
Biological characterisation of boron and BColl films in 2D: **a)** Metabolic activity and DNA content of osteoblasts in suspension with varying concentrations of boron nanoplatelets, showing robust proliferation, particularly for the highest boron concentration. **b)** Metabolic activity and DNA content of osteoblasts grown on BColl films, showing healthy proliferation in all conditions. **c)** Percentage coverage of osteoblasts grown on BColl films for 7 days, showing significant increase in coverage for cells grown on BColl films. **d)** Cell membrane (phalloidin) and nuclear (DAPI) staining of osteoblasts showed healthy cellular morphology and robust proliferation of cells grown on BColl films for 7 days. Imaged at 10× magnification. Significances: *p < 0.05, **p < 0.01, ***p < 0.001, ****p < 0.0001

### 2.5 Anti-microbial activity of BColl composite films

With the biocompatibility of the boron material confirmed in 2D, focus shifted to testing the anti-microbial activity of the BColl composite, since infection is a key concern during the implantation and lifetime of any scaffold. Escherichia coli (*E. coli*) and Staphylococcus aureus (*S. aureus*) were cultured on the surface of BColl films for 24 hours, followed by live-dead staining.

There are two primary origins of the anti-microbial effect caused by boron – first, boron compounds exhibit some innate anti-microbial effect with a range of bacterial strains.^[117–119]^ Second, 2D nanomaterials often exhibit anti-microbial behaviour due to the interaction of nanomaterial edges and basal planes with the bacterial cell membrane.^[120,121]^ There was a significant drop in *E. coli* viability at all concentrations of boron tested (**Figure 5**a). This indicates the potential of the boron to inhibit the growth of gram-negative bacteria on the surface of its composites, addressing a key clinical issue - Masters et al.^[122]^ states that 20% of all trauma-related skeletal infections and 35-55% of those in foot/ankle infections are caused by gram negative bacteria. However, it is important to note that no anti-microbial effect was observed with *S. aureus* bacteria (Figure 5b). This difference is potentially due to differing interactions of the boron edges with the bacterial membranes of gram-positive (*S. aureus*) vs. gram-negative (*E. coli*) bacteria. Gram-positive bacteria, while possessing more porous membranes and thus less resistance to typical anti-microbial agents, have considerably thicker cell membranes^[123]^ than gram-negative bacteria, which possess a thinner but more complex outer membrane. This thin membrane may be sliced more deeply by sharp nanoplatelets protruding from the composite surface, leading to a greater antimicrobial effect^[124–126]^ – however, further research will be required to establish the exact mechanism of the antimicrobial effect, especially in composite form. This indicates that boron nanoplatelets have the potential to be used in conjunction with a traditional anti-microbial agent such as chitosan or small-molecule antimicrobials, leading to effective inhibition of both bacterial subgroups across a range of strains, as demonstrated previously by Taşaltin et al.^[127]^ Furthermore, adaptations of the boron nanoplatelets, including size selection for smaller, thinner nanoplatelets, or inclusion of nano-needle (∼1D) or nano-sword (1D/2D) morphologies may improve the inhibitory effect of boron on bacterial growth.

**Figure 5.**
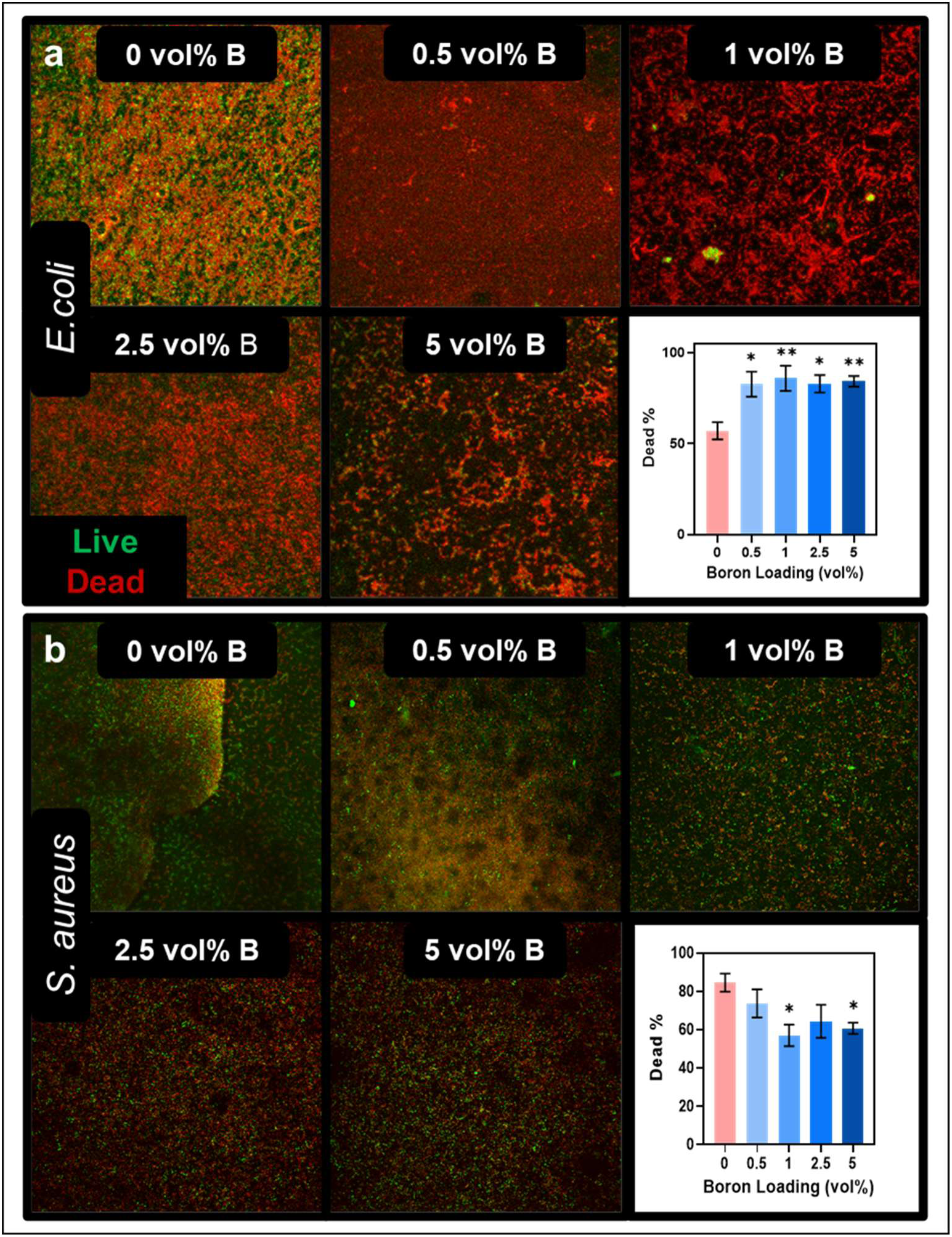
Anti-microbial testing of BColl films: Anti-microbial testing with BColl films of varying boron loadings ranging from 0–5 vol%, with E. coli **(a)** and S. aureus **(b)** bacteria grown for 24 h, showing significant anti-microbial activity against E. coli bacteria. Live-dead staining coverage was analysed using FIJI. Significances: *p < 0.05, **p < 0.01, ***p < 0.001, ****p < 0.0001

### 2.6 Biocompatibility and osteogenesis-inducing effect of BColl scaffolds with osteoblasts and mesenchymal stem cells (MSCs)

To demonstrate the potential of BColl in a 3D culture environment, osteoblasts were cultured in BColl scaffolds for 21 days. Both metabolic activity (**Figure 6**a) and DNA content analysis (Figure 6b) showed no significant difference from the control after 21 days, indicating a high degree of biocompatibility of BColl scaffolds with bone cells. The osteogenic potential of the boron was assessed by measuring the release of alkaline phosphatase (ALP), a key enzymatic regulator of bone formation,^[128]^ and by the deposition of calcium, indicative of bone mineralization.

**Figure 6.**
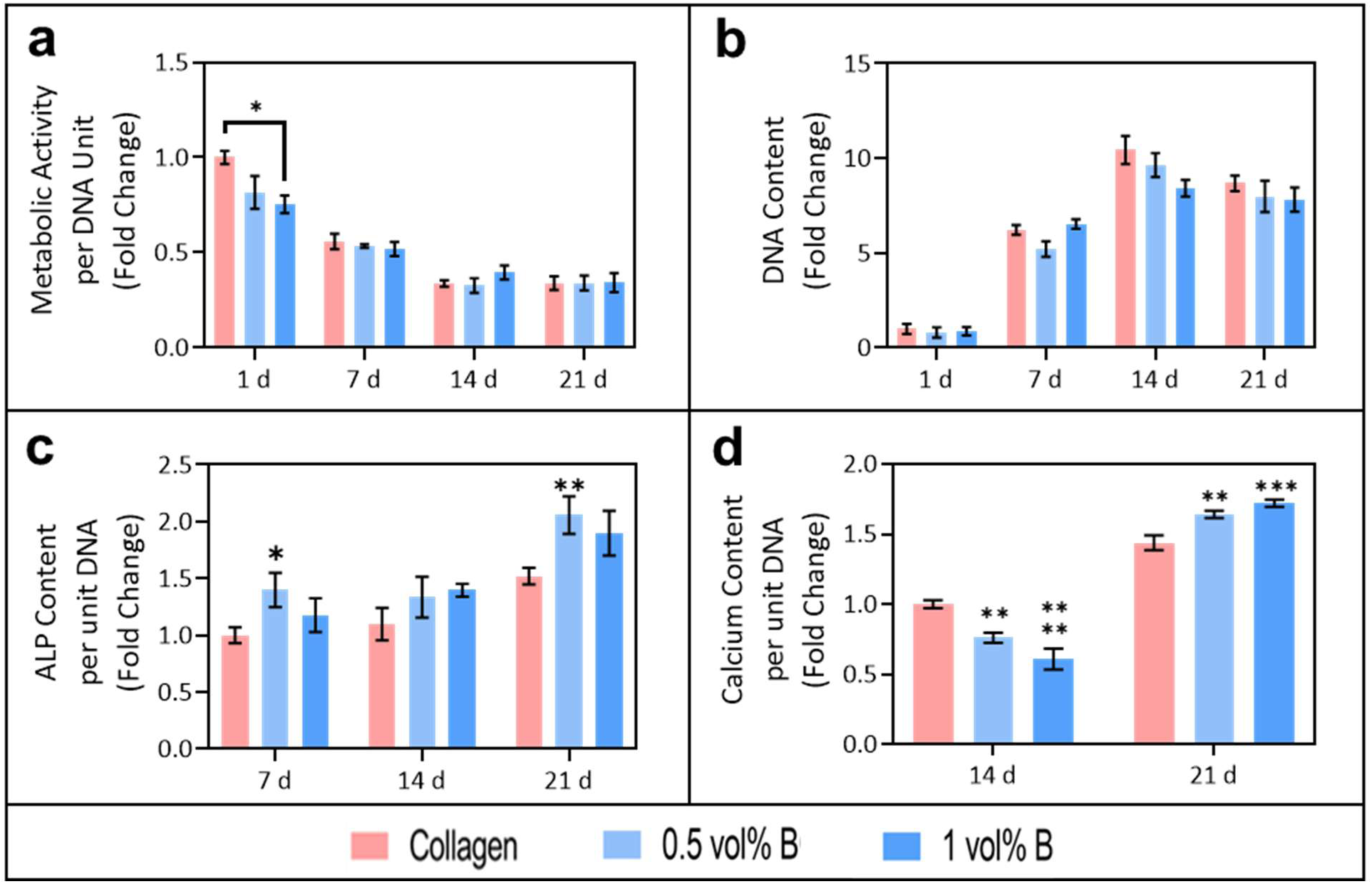
Assessment of osteogenic effect of BColl scaffolds with a bone cell line: a-b) Metabolic Activity (a) and DNA content (b) of osteoblasts grown on BColl scaffolds for 21 days, showing high levels of biocompatibility. **c)** ALP release of osteoblasts on BColl scaffolds, trending towards increased release for BColl scaffolds. **d)** Calcium release of osteoblasts on BColl scaffolds, showing a significant increase indicative of increased bone mineralization on BColl scaffolds. Significances: *p < 0.05, **p < 0.01, ***p < 0.001, ****p < 0.0001

There was a trend towards increased ALP release for the BColl scaffolds at each measured timepoint (Figure 6c) and calcium deposition showed a significant increase at 21 days (Figure 6d), indicative of improved bone mineralization. This shows that boron supplementation leads to robust proliferation and enhanced osteogenic behaviour in osteoblasts.

While the MC3T3 osteoblast cell line is a good model for osteoblast mineralization in general, they are not perfect models, with a differing gene expression profile, and variable degrees of mineralization when compared with the primary osteoblasts they are intended to replicate.^[129]^ For these reasons, we decided to assess the biocompatibility, osteogenic potential, and functional performance of the boron nanomaterial with a more physiologically relevant cell type. For this purpose, mesenchymal stem cells (MSCs) are highly therapeutically relevant, as they are readily differentiated to an osteoblastic lineage by media supplementation and are used clinically in musculoskeletal applications.^[130,131]^ By culturing MSCs on BColl scaffolds, we are thus able to more accurately model long-term clinical implantation. MSCs were grown on BColl scaffolds for 28 days followed by fluorescent staining and confocal imaging, showing that the cells had proliferated robustly across the BColl scaffolds, and maintained a healthy cellular morphology (**Figure 7**a). and the biocompatibility was assessed by metabolic activity (Figure 7b) and DNA content (Figure 7c) analysis.

**Figure 7.**
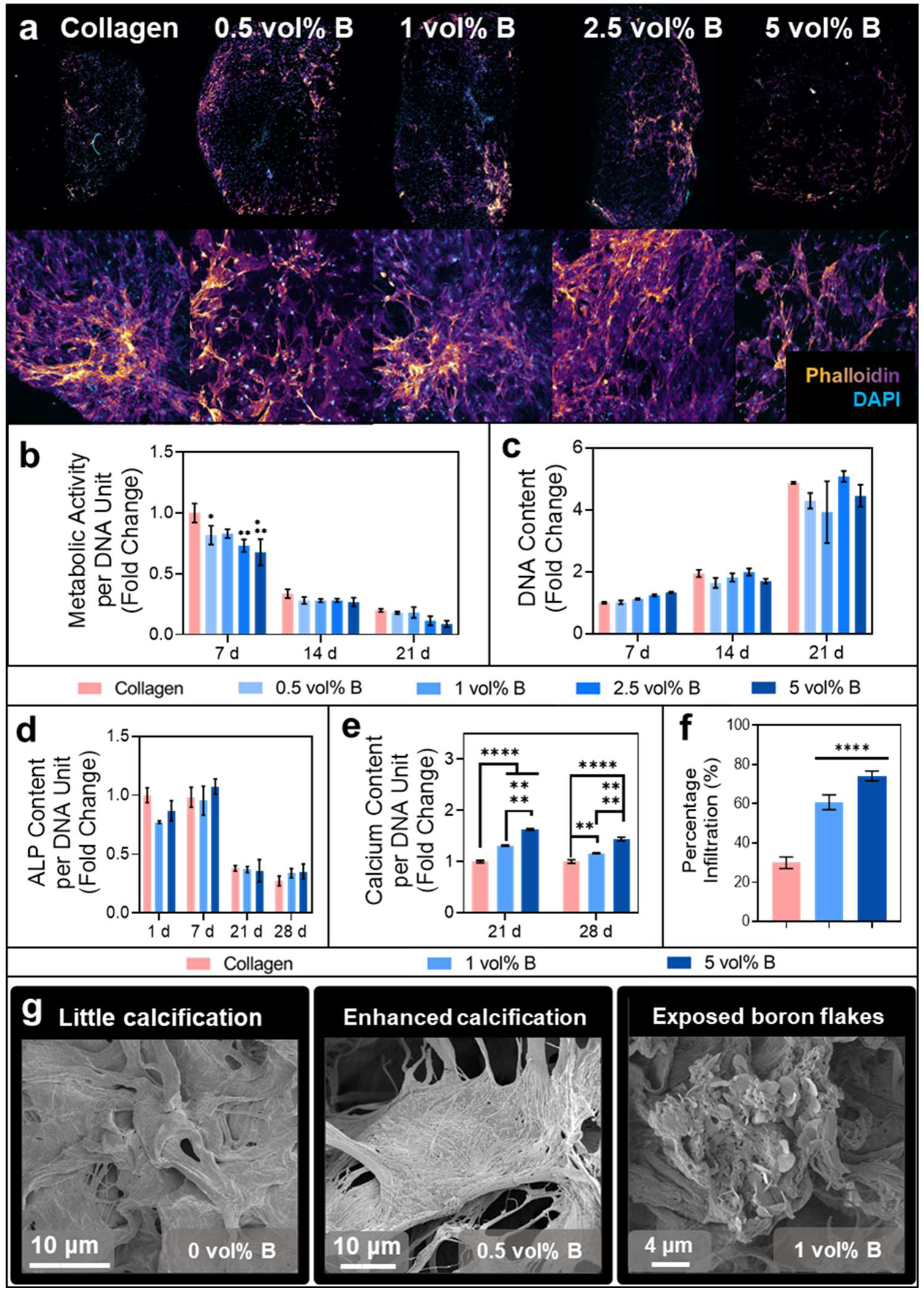
Assessment of osteogenic effect of BColl scaffolds with osteogenically-differentiated mesenchymal stem cells: **a)** Confocal microscope images of lateral cross sections of BColl scaffolds with MSCs grown in them for 21 days, stained with phalloidin for the cell membrane, and DAPI for the cell nucleus, showing robust cellular proliferation and infiltration of the scaffolds. **b)** Metabolic activity of MSCs on BColl scaffolds and **c)** DNA content of MSCs on BColl scaffolds indicate high levels of biocompatibility. **d)** ALP release of MSCs on BColl scaffolds, showing significant fall-off in release after differentiation is complete. **e)** Calcium release of MSCs on BColl scaffolds, showing significant, dose- dependent increase in calcium deposition and thus bone mineralization. **f)** Percentage infiltration of MSCs into boron scaffolds measured radially from the geometric center of the scaffold, showing significant increase in infiltration for boron-containing scaffolds. **g)** SEM imaging showing increased calcification in BColl scaffolds, and exposed boron flakes in the collagen matrix. Significances: *p < 0.05, **p < 0.01, ***p < 0.001, ****p < 0.0001

No significant difference was observed between the boron-treated samples and the collagen-only controls, indicating that the BColl material is well-tolerated even by sensitive MSCs at high concentrations, up to 5 vol%, for extended periods of culture. The osteogenic potential of these cells was then assessed, as before, by analyzing ALP release and measuring calcium deposition. There was no significant difference observed in the ALP release for the BColl samples (Figure 7d) - however, ALP release has a short peak, typically around day 8 as the differentiating cells commit to the osteoblastic lineage,^[132]^ and thus it may have been missed by the timepoints analysed. This peak is reflected in the higher per cell expression of ALP for all conditions at days 1 and 7, compared to the later timepoints. As described above, the key indicator of osteogenic potential of any biomaterial is the quantity of calcium deposition. This was measured at days 21 and 28, when MSC-derived osteoblasts had matured sufficiently to lead to calcium deposition (Figure 7e). A significant dose-dependent increase was seen in the calcium deposition for BColl samples when compared with collagen-only scaffolds. This indicates that boron is having a positive effect on the rate of mineralization, and thus the osteogenic potential of the cells, pointing towards the potential of BColl scaffolds to accelerate bone formation after implantation. Cross-sections of these scaffolds were stained, and infiltration analysis showed that cells in BColl scaffolds were colonizing deeper into the scaffold (Figure 7f). Finally, SEM (Figure 7g) and EDX (**Figure S7**) of these scaffolds showed increased calcium phosphate deposition for boron containing scaffolds, and further SEM showed areas with exposed boron flakes on the higher loading samples (Figure 7g). While the increase in calcium deposition with boron scaffolds is lower compared to additives like osteoconductive hydroxyapatite^[133]^ or expensive osteoinductive growth factors BMP-2,^[134]^ these scaffolds exhibit both osteoconductive and osteoinductive properties. Additionally, boron scaffolds offer a multifunctional profile, encompassing osteogenic, angiogenic, neurogenic, anti-microbial, mechanically reinforcing, and anti-inflammatory effects that address key challenges in bone repair. Achieving such comprehensive functionality would otherwise require the use of multiple, potentially incompatible additives. Furthermore, although growth factors like BMP-2 provide strong osteoinductive effects, they are associated with several adverse outcomes at high concentrations and high cost when used clinically.^[135]^

### 2.7 Neurogenic assessment of BColl scaffolds with dorsal root ganglia

To assess the impact of boron loading on innervation, a key functional outcome in bone repair, the neurogenic effect of BColl was assessed. Dorsal root ganglia (DRGs) isolated from rats, a model of neuronal tissue that is both aged and injured, and thus particularly difficult to heal, were cultured on the surface of BColl scaffolds for 14 days. Staining and confocal imaging subsequently established that DRGs extended neurites effectively on BColl scaffolds and maintained healthy morphologies (**Figure 8**a). We next established that the scaffolds led to no cytotoxic or inflammatory effect, by measuring LDH (Figure 8b) and pro-inflammatory TNF-α release (Figure 8c), which showed no significant difference compared to the collagen control scaffolds.

**Figure 8.**
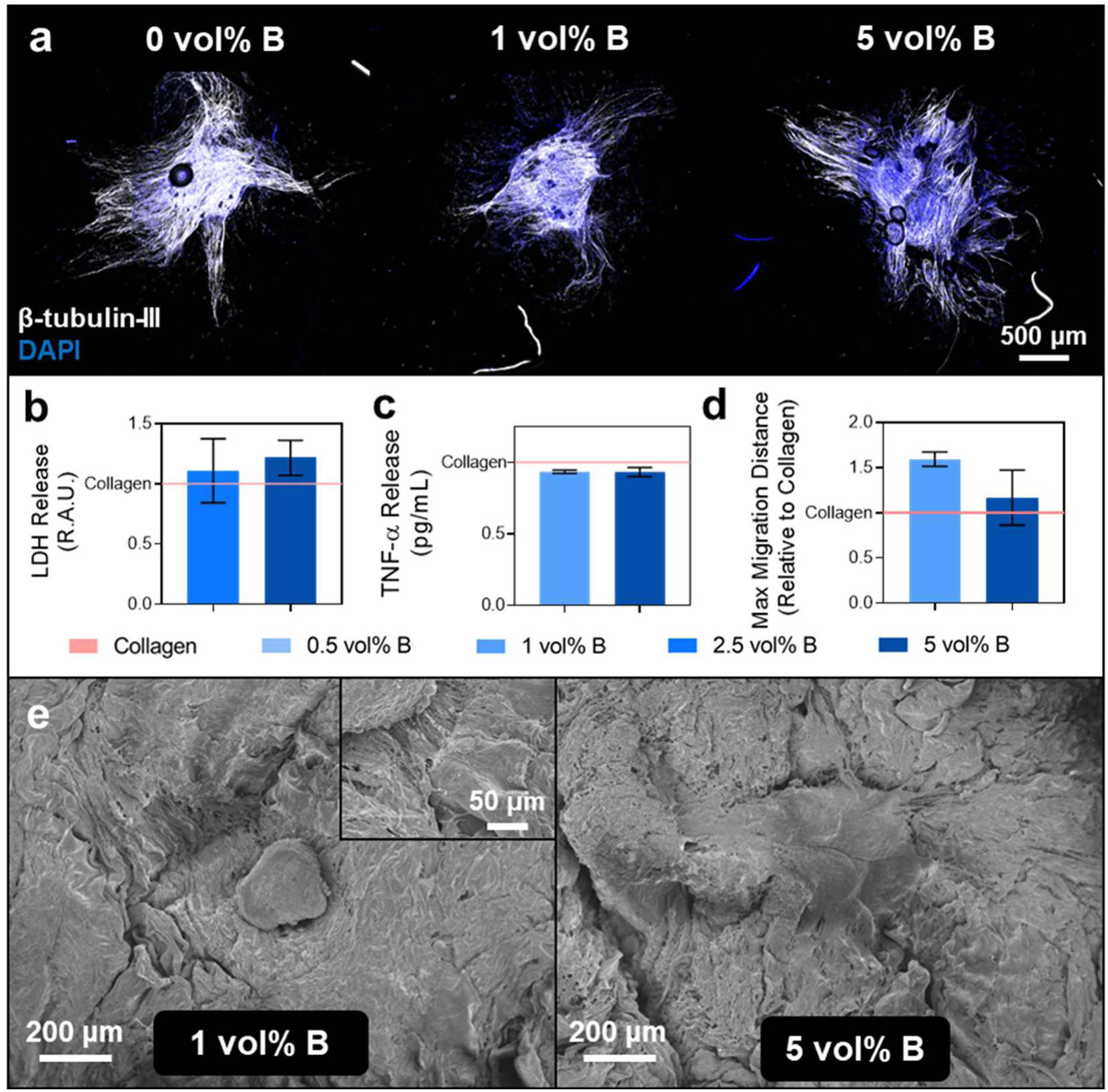
Assessment of neurogenic effect of BColl scaffolds on dorsal root ganglia (DRGs): ***a)*** Representative confocal microscopy images of DRGs on surface of scaffolds, stained with β-III-tubulin as a marker of neuronal maturity and for cell morphology, and counter-stained with DAPI for nuclei. ***b)*** Lactate dehydrogenase (LDH) release was assessed to investigate the cytotoxicity of BColl scaffolds. ***c)*** Tumor Necrosis Factor alpha (TNF-α) release was assessed to examine the effect of BColl scaffolds on the inflammatory signaling response of DRGs, showing a slight trend towards decreased TNF-α release. ***d)*** Maximum migration distance of neurites from DRG center showing a trend towards increased migration for BColl scaffolds, measured from confocal images by using NeuriteTracer^[136]^ and FIJI. **e)** SEM images of DRGs growing on BColl scaffolds, with significant neurite outgrowth visible on both samples

The neurite outgrowth from the DRGs was extracted from these images, showing an increase in the maximum migration distance of DRG neurites on BColl scaffolds (Figure 8d). This was qualitatively confirmed by SEM imaging of the same scaffolds, showing significant neurite outgrowth on BColl scaffolds (Figure 8e). This data shows that boron contributes to a supportive environment for neurogenesis and innervation, a key process in bone regeneration,^[137,138]^ by employing a highly physiologically relevant tissue explant model.

### 2.8 Angiogenic, immunomodulatory, and neurogenic effect of BColl scaffolds on MSCs

The final aim of this study was to demonstrate the benefits of boron incorporation on the paracrine release profile of mesenchymal stem cells, a key cell type that modulates bone remodeling *via* cross-talk with other bone cells, and *via* their differentiation into osteoblasts.^[139]^ Effects on the paracrine signaling of these cells can potentially have a downstream impact on the therapeutic efficacy of the scaffold, so a multiplex ELISA array was used to assess the cytokine release profile of mesenchymal stem cells on the BColl scaffolds at day 7 (**Figure 9**a). Of course, many of the cytokines assessed using this array have multiple potential effects, which were assessed by a literature review (Figure 9b). Firstly, several cytokines can be used to corroborate the calcium deposition data, indicative of enhanced osteogenic signaling and a more favorable environment for osteoblastic differentiation. Several factors involved in both osteogenesis (monocyte chemoattractant protein-1 (MCP-1 (CCL2)), macrophage inflammatory protein-3 alpha (MIP-3α (CCL20))^[140]^) and osteoclastogenesis (interleukin-1α/β (IL-1α/β),^[141]^ fractalkine (CX3CL1)^[142]^ and tumor necrosis factor alpha (TNF-α)) were upregulated in BColl scaffolds. We posit that the apparent dose-dependent increase in these osteogenic and osteoclastogenic factors results in a more favorable scaffold environment for infiltrating MSCs, which in turn impacts the rate of osteoblastic differentiation and bone remodeling and deposition within the scaffold. It is well-established that an increase in osteoblast activity will often be associated with an increase in osteoclast activity, to maintain bone homeostasis.^[143]^

**Figure 9.**
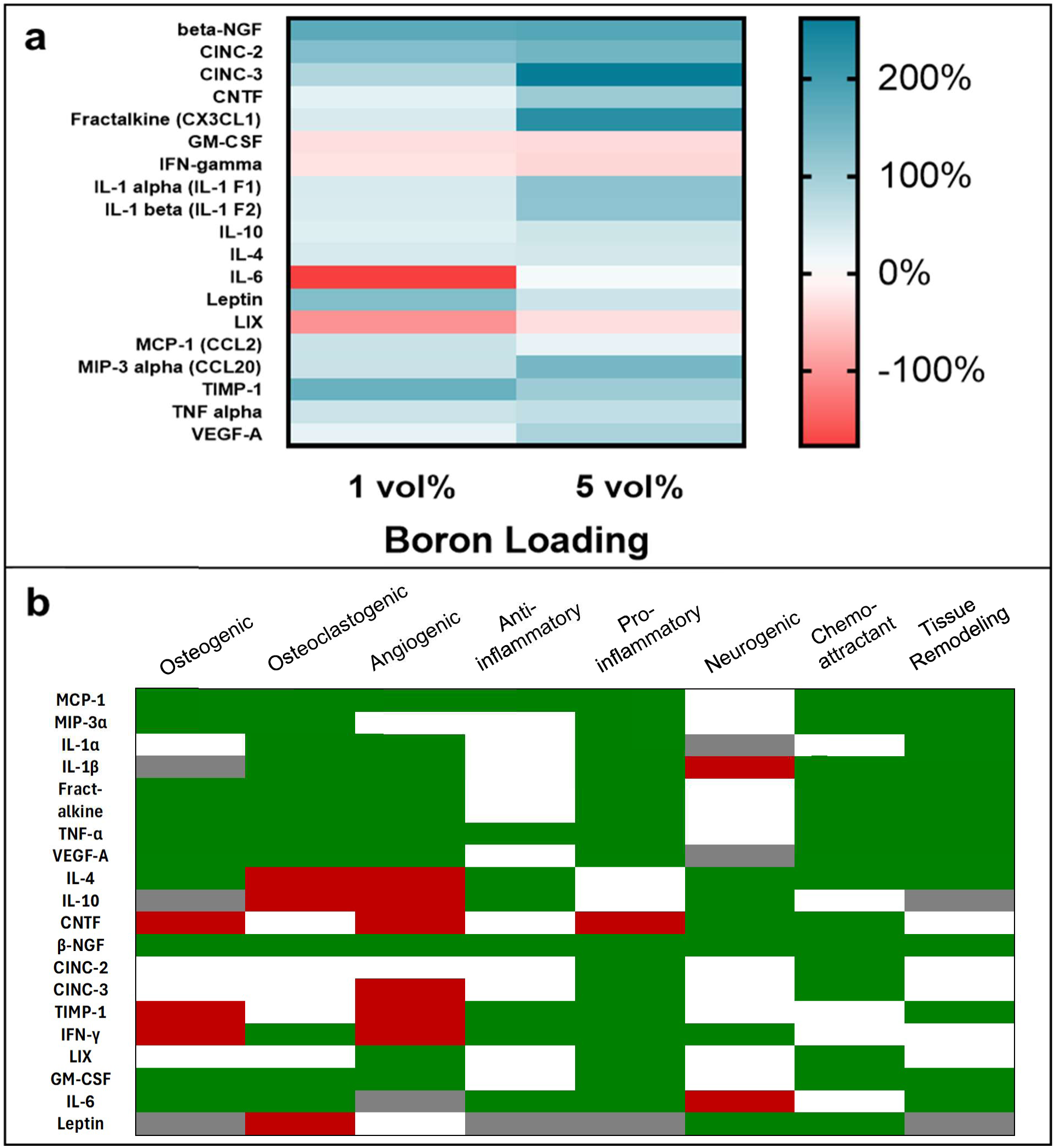
Investigation of paracrine release profile of rMSCs on BColl scaffolds: **a)** Assessment of cytokine release profile of rat mesenchymal stem cells using a multiplex ELISA array. Heatmap indicating relative change to collagen-only control. **b)** Matrix detailing the primary effects of each cytokine assessed in the ELISA data. Green corresponds to literature evidence of an effect, grey to contradictory or highly environment-dependent effect, red corresponds to evidence of a negative effect and white corresponds to no evidence for an effect. Citations are available in **Figure S10**

Tissue inhibitor of metalloproteinase 1 (TIMP-1), a cytokine involved in tissue remodeling, is also upregulated in the BColl scaffolds, corroborating the shift towards a supportive environment for bone repair. A dose-dependent increase was observed in the expression of angiogenic factors such as vascular endothelial growth factor A (VEGF-A), fractalkine^[144]^ and lipopolysaccharide-induced CXC chemokine (LIX).^[145]^ This effect has been observed in boron-containing compounds in the literature, with increased phosphorylation in the Akt/eNOS pathway proposed as the mechanism for its upregulatory effect.^[146]^ The pro-angiogenic effect is also backed up by a strong trend for enhanced VEGF release from a human ELISA assay on the BColl samples (**Figure S11**). This is a key finding, as vascularization is an important target in bone regenerating biomaterials,^[147]^ and the induction of infiltrating MSCs to a pro-angiogenic phenotype renders the scaffold a more favorable environment for blood vessel formation. Enhanced vascularization is not the only non-bone effect essential for bone repair however – innervation is required for healthy bone repair.^[79]^ Both nerve growth factor-beta (β-NGF) and ciliary neurotrophic factor (CNTF) were upregulated in the BColl scaffolds, indicating an improvement of the neurogenic environment of the scaffold following boron addition, and corroborating the results found with dorsal root ganglia. This is consistent with the literature, where boron-containing compounds have been previously evaluated for their neuroprotective and neuroregenerative effects.^[137]^ Finally, increased anti-inflammatory signaling of MSCs in BColl scaffolds was indicated by downregulation of key inflammatory cytokines interleukin-6 (IL-6), interferon gamma (IFN-γ) and granulocyte-macrophage colony-stimulating factor (GM-CSF), with associated upregulation of anti-inflammatory cytokines such as interleukin-4 (IL-4) and interleukin-10 (IL-10). This is a highly notable result, as nanomaterials often cause inflammatory responses in biological systems.^[148]^ This anti-inflammatory effect of boron has been observed in previous work,^[17,149,150]^ suggesting that boron-mediated immunomodulation may be countering any nanomaterial-induced inflammatory effect. It is worth addressing that many of the cytokines implicated in osteogenesis, osteoclastogenesis, and angiogenesis above are also associated with inflammatory and immune responses from macrophages and other cell types *in vivo*. However, taking the data as a whole, it is clear that boron incorporation in the scaffolds leads to healthy, proliferative cells with a pro-osteogenic paracrine signaling profile. This, coupled with the downregulation of key pro-inflammatory cytokines, makes it significantly more likely that the upregulation of these pleiotropic cytokines is associated with reparative processes.

## 3 Conclusion

This study demonstrates that boron nanoplatelets can be used as a versatile and powerful additive for enhancing the regenerative properties of biomaterial scaffolds for bone repair, thanks to their unique combination of intrinsic and 2D material-specific properties. High aspect ratio 2D boron nanoplatelets, stabilized with polyvinylpyrrolidone, were synthesized using an optimized liquid-phase exfoliation approach. Their biocompatibility was confirmed across a range of exposure conditions and cell types, demonstrating their safety for biomedical applications. Boron increased the osteogenic potential of both osteoblasts and osteogenically differentiated mesenchymal stem cells in long-term 3D scaffold culture. The material also boosted angiogenic, neurogenic and anti-inflammatory cytokine signaling, key targets for regenerative therapies, in addition to increasing neurite migration of neuronal tissue explants on scaffolds. Additionally, the 2D morphology of the nanoplatelets yielded anti-microbial activity against *E. coli*, and significantly enhanced the tensile modulus, toughness, and strength of collagen. These findings highlight BColl as a versatile composite biomaterial, offering multifunctional benefits for key bone regeneration targets such as mineralization, angiogenesis, and innervation, along with improved mechanical and antimicrobial properties. In summary, this study employs cutting-edge nanoscience and novel exfoliation techniques to develop biocompatible boron nanoplatelets and BColl composite scaffolds, which represent a promising new treatment for tackling bone repair in orthopedic, reconstructive, and tissue engineering applications.

## 4 Methods

### 4.1 Synthesis of 2D boron nanoplatelets

#### 4.1.1 Preparation of inert solvents to prevent boron oxidation

Inert, anhydrous, and deoxygenated solvents were produced by drying, distillation and degassing. The solvent was first dried using molecular sieves for 3 days. The solvent was then distilled to remove impurities. Finally, it was degassed by bubbling nitrogen gas through it for 2 days. The material was then stored in the glovebox under an argon atmosphere until use.

#### 4.1.2 Sonic tip exfoliation of boron nanoplatelets

Crystalline boron powder (Stanford Advanced Materials, median particle diameter = 15 µm), stored under an argon atmosphere, and 160 mg of polyvinylpyrrolidone (PVP, Mw = 40 kDa, Sigma Aldrich) were added to 80 mL of anhydrous, degassed *N*-Methyl-2-pyrrolidone (NMP, ACS Reagent, ≥99% purity, Honeywell, USA) in a metal beaker. The mixture was then exfoliated for 8 h at 55% amplitude, on a 6 s/2 s on/off pulse, using a sonic probe (VCX750, Sonics & Materials Inc., USA). Following exfoliation, the material was centrifuged at 1 krpm to cause unexfoliated material to crash out of the dispersion. This sediment was discarded, and the supernatant was subsequently centrifuged at 6 krpm, to remove small impurities and 0D particles from the dispersion.^[151]^ The resulting sediment was redispersed in anhydrous, degassed isopropanol (IPA, HPLC grade, 99.9% purity, Sigma Aldrich), and washed twice with fresh IPA at 6 krpm to remove excess residual NMP. To slow oxidation, the resulting dispersion was stored under argon in a glovebox (Jacomex, France) until use. For use in biological scaffolds, the material was redispersed in deionized water (18.2 MΩ) with 2 mg mL^-1^ polyvinylpyrrolidone (PVP, Sigma) by centrifugation at 6 krpm, and washed twice in fresh deionized water to remove traces of IPA. The concentration of the dispersions was assessed by filtration of a known volume of dispersion using a 200 nm filter membrane and weighing.

#### 4.1.3 Cryogenic exfoliation of boron

Cryogenic exfoliation was carried out by submerging a vial of 1 mg mL^-1^ boron powder and 1 mg mL^-1^ PVP in IPA in liquid nitrogen for 5 minutes. The material was then immediately placed in a sonic bath and sonicated at 37 kHz for 1 hour.

### 4.2 Physical characterisation of boron nanoplatelets

#### 4.2.1 Establishing thermal degradation and oxidation properties using thermogravimetric analysis (TGA)

Thermogravimetric analysis was carried out using a Q50 thermogravimetric analyzer (TA Instruments, USA). The temperature was ramped from 25 °C to 900 °C, at a ramp rate of 10 °C min^−1^ in air, then held isothermal at 900 °C for 5 min to ensure complete combustion.

#### 4.2.2 Elemental and oxidation analysis using energy-dispersive X-ray spectroscopy (EDX)

EDX analysis was carried out using the secondary electron detector of a Zeiss Ultra FE-SEM (Zeiss, USA), at an accelerating voltage of 5 kV and a working distance of 11 mm. Peak assignment was carried out using INCA software (Oxford Instruments).

#### 4.2.3 Scanning electron microscopy (SEM) for nanoplatelet and scaffold visualization

Where necessary, samples were first gently dehydrated using a Leica critical point dryer. Dry samples were then sputter coated with a 5 nm layer of an 80:20 mixture of gold and palladium using a Cressington 108 auto sputter coater. Imaging was carried out using a Zeiss Ultra FE-SEM using the InLens detector at an accelerating voltage of 5 kV, an aperture size of 30 μm and a working distance of 5 mm, with images acquired at varying magnifications.

#### 4.2.4 Ultraviolet-visible (UV-Vis) spectroscopy

Boron nanoplatelets were first diluted 1:20 in isopropanol. UV-Vis spectroscopy was then carried out in a quartz cuvette (path length 1 mm) using a Perkin-Elmer Lambda 1050+ spectrophotometer and an integrating sphere, to collect information on the scattering, absorption, and extinction spectra. A background spectrum with isopropanol was gathered as a reference and subtracted from all spectra.

#### 4.2.5 Atomic force microscopy (AFM) for establishing nanoplatelet dimensions and morphology

Utilizing a Bruker Multimode 8 microscope, AFM was performed to assess the thickness and lateral dimensions of the flakes. After dilution with isopropanol at a ratio of 1:100, the boron inks were drop-cast onto Si/SiO2 substrates. Imaging of the samples was carried out using OLTESPA R3 cantilevers in ScanAsyst mode, and statistical data were derived from the analysis of 569 flakes. The lateral dimensions were determined by computing the square root of the product of the flake length and width. Data was analysed using Gwyddion SPM software.^[152]^

#### 4.2.6 Transmission electron microscopy (TEM) for nanoplatelet visualization

A JEOL 2100 system operating in bright-field mode was used to perform the transmission electron microscopy measurements. Freshly exfoliated boron nanoplatelets, centrifuged between 1 and 6 krpm, were used to prepare the holey carbon TEM grids (400 mesh). 100 µL of the dispersion (concentration of approx. 0.1 mg mL^-1^) was drop-casted on the grids, and vacuum dried at 80 °C in the oven overnight prior to the measurements. Images were acquired at an accelerating voltage of 200 kV.

#### 4.2.7 X-ray diffraction (XRD) for establishing material crystallinity

The powder samples were structurally characterized by performing symmetric 2*θω* X-ray diffraction (XRD) scans using a Panalytical X’Pert Pro diffractometer equipped with a Cu K*α* source (*λ* = 1.5406 Å) and a Ni filter to remove K*β* peak intensities.

### 4.3 Mechanical testing to identify mechanical reinforcement of natural polymers

Mechanical testing was carried out using a Zwick Z0.5 Proline tensile tester with a 100 N load cell, at a strain rate of 0.01 %*/*s. Samples were cut into strips (20 mm × 3 mm), clamped, and tested in a uniaxial strain configuration. The Young’s modulus and other mechanical properties were calculated for each sample from the stress-strain curve.

### 4.4 Synthesis of 2D boron/collagen (BColl) nanocomposites

The BColl slurry was fabricated by mixing type I collagen (bovine tendon, 5 mg mL^-1^, milled form, Collagen Solutions) with a 2D boron dispersion in deionized water and acetic acid. The composite slurry was blended (T25 Ultra-Turrax®, VWR) at 15000 rpm for 30 min until homogenous. The composite materials subsequently underwent dehydrothermal (DHT) crosslinking treatment for 24 h at 105 °C under vacuum. For 2D culture, they were cast in square molds and allowed to dry. For 3D culture, porous scaffolds of BColl were produced by lyophilization - slurries were placed in Teflon molds and freeze dried for 24 h at -40 °C to obtain lyophilized cylindrical scaffolds.

### 4.5 Biocompatibility testing of 2D boron sheets

#### 4.5.1 Ethical declaration for ex vivo experiments

For the experiments detailed in Sections 4.5.2 and 4.8, *ex vivo* tissue was obtained from animals donated as post-mortem byproducts from ongoing experiments in other groups in the RCSI, in line with our policy of reduction according to the 3Rs of animal research. Rat femora for isolating mesenchymal stem cells in Section 4.5.2 were obtained from rats culled under project authorisation number REC202012003, accredited by the Animal Research Ethics Committee of RCSI (RECTH017). Mouse spinal cords used to extract dorsal root ganglia in Section 4.8 were obtained from animals donated by the Tissue Engineering Research Group. Post-mortem harvesting was carried out under Health Products Regulatory Authority (HPRA) individual license AE19127/I259, with ethical approval from the Animal Research Ethics Committee of RCSI (RECTH017).

#### 4.5.2 Isolation of clinically relevant rat mesenchymal stem cells (rMSCs)

Rat mesenchymal stem and endothelial progenitor cells were isolated from 6–8-week-old male Sprague Dawley rats as described in Power *et al*.^[153]^, under ethical approval detailed in Section 4.5.1. Both ends of the femora of the hind legs were clipped to expose the bone marrow, which was then flushed out with rat MSC medium using an 18 G needle and a 12 mL syringe into a petri dish using Dulbecco’s Modified Eagle Medium (DMEM) containing: 20% fetal bovine serum (FBS, Biosera, Ireland), 2% penicillin streptomycin (P/s, Sigma Aldrich, USA), 1% non-essential amino acids (NEAA, Thermo Fisher Scientific, USA), 1% GlutaMAX (Thermo Fisher Scientific, USA) and 0.002% Primocin (Sigma, Ireland). The cells were then cultured in 15 mL of media under standard cell culture conditions (37 °C, 5% CO2) for 24 h. The supernatant was then removed and centrifuged at 300 × g for 5 min.

#### 4.5.3 2D culture of osteoblasts in free suspension with boron nanoplatelets

2D boron was first dispersed in DI water and sterilized by autoclaving. Osteoblastic immortalized cells (MC3T3-E1, isolated from mouse calvaria, American Type Culture Collection, ATCC, Middlesex, UK) were cultured in T175 flasks until needed, then seeded in a 48-well plate, at a density of 1×10^4^ cells per well. The cells were supported in growth media consisting of Alpha Modified Eagle Medium (α-MEM, Sigma, Ireland) with 10% fetal bovine serum (FBS, Biosera, Ireland), 1% Penicillin/Streptomycin (P/s, ScienCell, USA) and 1% l-glutamine (l-glut, Sigma, Ireland). The cells were seeded, and left to settle for 24 h, before the treatment was applied. Dispersions of 40 µg mL^-1^ and 80 µg mL^-1^ of boron nanoplatelets in growth media were made up, and the media on the cells was replaced with 360 µL of fresh treatment media after 24 h of settling. After 24 h and 72 h, 40 µL of Alamar Blue (Invitrogen, UK) was added to measure the cellular metabolic activity, and the cells were incubated at 37 °C for 2 h. Following this, fluorescence was measured using excitation 535 nm and emission 590 nm with a plate reader (Infinite M Plex, Tecan), to assess the metabolic activity of the cells. The cells were then lysed using lysis buffer (100 mL autoclaved DI water, 0.5 mL Triton X- 100, 29 mg EDTA (Ethylenediaminetetraacetic acid), 2.12 g sodium carbonate) and total DNA content was assessed using the Picogreen assay (Invitrogen, UK), which was carried out as per the manufacturer’s instructions. For assessment of osteogenesis in 3D, 1% β-glycerophosphate (Sigma, Ireland), 0.5% ascorbic acid (Sigma, Ireland) and 100 nM dexamethasone (Sigma, Ireland) were added to create osteogenic differentiation media.

#### 4.5.4 2D culture of osteoblasts on BColl films

BColl films were fabricated as described above, at loadings of 0 vol%, 0.5 vol%, and 1 vol%. MC3T3 cells were seeded on 8 mm diameter discs of these films and allowed to grow for 7 days.

#### 4.5.5 3D culture of osteoblasts and MSCs on BColl scaffolds

BColl scaffolds were fabricated as described above, at loadings of 0 vol%, 0.5 vol%, 1 vol%, 2.5 vol%, and 5 vol%. Scaffolds were sterilized in 70% ethanol for 24 h, followed by washing in PBS (×3, 5 mins). The scaffolds were then immersed in growth media at 37 °C and 5% CO2 for 24 h. Cells were passaged and seeded at densities of 3×10^5^ – 5×10^5^ per scaffold. Half of the cells were seeded on each side of the scaffold for 1 hour each, before the wells were flooded with growth media for 24 h. Growth media was exchanged for osteogenic media after 24 hours.

### 4.6 Analysis of boron effect on osteogenesis and bone mineralization

#### 4.6.1 Alkaline phosphatase (ALP) release quantification to assess osteogenesis

ALP release was analysed using a Sensolyte™ PNPP ALP Assay Kit (Anaspec, USA). Cells were washed with PBS at the relevant timepoints, and subsequently lysed with ALP lysis buffer. The assay was then carried out according to the manufacturer’s instructions.

#### 4.6.2 Calcium (Ca^2+^) release quantification to assess bone mineralization

Following culture, calcium was extracted from quartered scaffolds using 0.5 M hydrochloric acid (Sigma, Ireland). Calcium release was analysed using a Calcium LiquiColor™ assay (StanBio, USA), according to the manufacturer’s instructions.

#### 4.6.3 Histology

##### Sample Preparation

Samples were embedded in sucrose for 48 h, and then sectioned in OCT using a cryostat (Leica Biosystems, UK), into 5 µm thick slices.

##### Hematoxylin & Eosin (H&E) Staining

H&E staining was carried out to image the ECM and nuclei present in the sample. The following order was used for staining: Tap water rinse for 5 minutes, hematoxylin for 3 minutes, tap water for 10 minutes, DI water for 5 minutes, eosin for 3 minutes, 3-4 dips in tap water, 15 dips in 95% ethanol, 15 dips 100% ethanol, 15 dips in acetone and 15 dips in xylene substitute. Coverslips were then mounted on the samples using DPX.

##### Alizarin Red Staining

Alizarin Red staining was carried out to image calcium deposition present in the sample. The following order was used for staining: Tap water rinse for 5 minutes, 250 µL of 2% Alizarin Red (Generon, Ireland) per slice for 30 minutes, 5 dips in acetone, 20 dips in 1:1 acetone:xylene substitute solution, xylene substitute for 5 minutes. Coverslips were then mounted on the samples using DPX.

#### 4.6.4 Immunofluorescence Staining

Cells were fixed using 10% formalin (Sigma, Ireland), and stained with phalloidin (Sigma, USA) for the cell membrane, and DAPI (Thermo-Fisher Scientific, USA) for the cell nucleus. The samples, once fixed and stained, were fluorescently imaged using a Zeiss AxioObserver inverted microscope. Infiltration analysis was carried out by measuring the radial distance from the edge of the scaffold to the furthest infiltrated cell at 15° intervals around the cross-section.

### 4.7 Anti-microbial testing

**5** *S. aureus* (Newman) and *E. coli* bacteria were cultured separately for 24 h at 37 °C in brain/heart infusion broth, before 1 mL of bacteria was plated into wells containing sterile boron discs (diameter = 6 mm) at a density of 5 × 10^5^ CFUs mL^-1^. These samples were then cultured for another 24 h at 37 °C. The broth was subsequently removed, and the samples were washed with phosphate buffered saline. Live-dead staining was then carried out with SYTO 9 and propidium iodide (LIVE/DEAD® BacLight™ Bacterial Viability Kit L7007, Thermo Fisher Scientific, USA), according to manufacturer instructions. Imaging was carried out on a Zeiss LSM 710 NLO confocal microscope and analysed using FIJI.

### 4.8 DRG culture with BColl scaffolds

To assess the ability of the scaffolds to enhance neurogenesis in an ex vivo model, dorsal root ganglia (DRGs) from 5-month-old adult female C57 mice (N=9, 385 – 500 g) were seeded onto the scaffolds. All animals were kindly donated by the Tissue Engineering Research Group, under the ethical approval detailed in Section 4.5.1. DRGs were isolated and dissected using a previously established method^[102]^. 20-24 DRGs and their roots were dissected from each animal. The roots were then trimmed of their associated nerves using micro-scissors (F.S.T. Cat # 15000-08) and cultured on scaffolds in neurobasal medium with 1% P/s, 1% GlutaMAX and 2% B27 supplement for 14 days.

### 4.9 ELISA Analysis

ELISA analysis was carried out using a 19-target multiplex array of rat cytokines (AAR-CYT-1-4, CliniSciences, Ireland). Samples were diluted 1:2 with blocking buffer, and analysis was carried out according to manufacturer’s instructions. Densitometry measurements were carried out using FIJI. A blank reading consisting of cell-free media was subtracted from the data.

## 5 Acknowledgements

Additionally, we would like to acknowledge the support given by the Advanced Microscopy Laboratory (AML, Trinity College Dublin) in the acquisition of SEM data.

## 6 Data Availability Statement

The data that support the findings of this study are available from the corresponding author upon reasonable request.

## Supporting Information

**Fig. S1.**
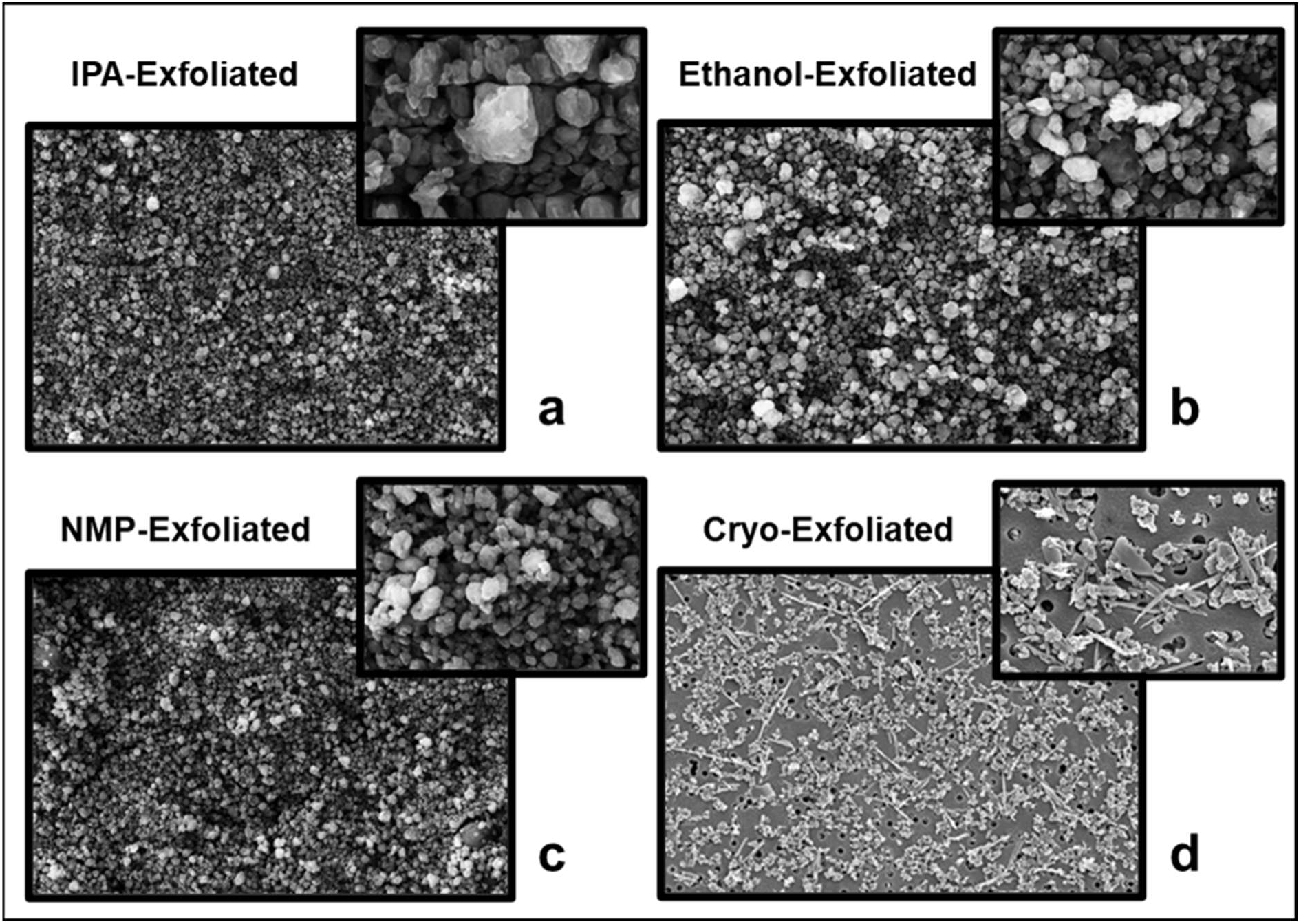
Exfoliation of boron in various solvents: a-c) Exfoliation of amorphous boron using a sonic tip in isopropyl alcohol (a), ethanol (b) and NMP (c), leading to nanoparticulate morphology. **d)** Cryogenically assisted exfoliation of amorphous boron in NMP, leading to a mix of 0D, 1D and 2D morphologies

**Fig. S2.**
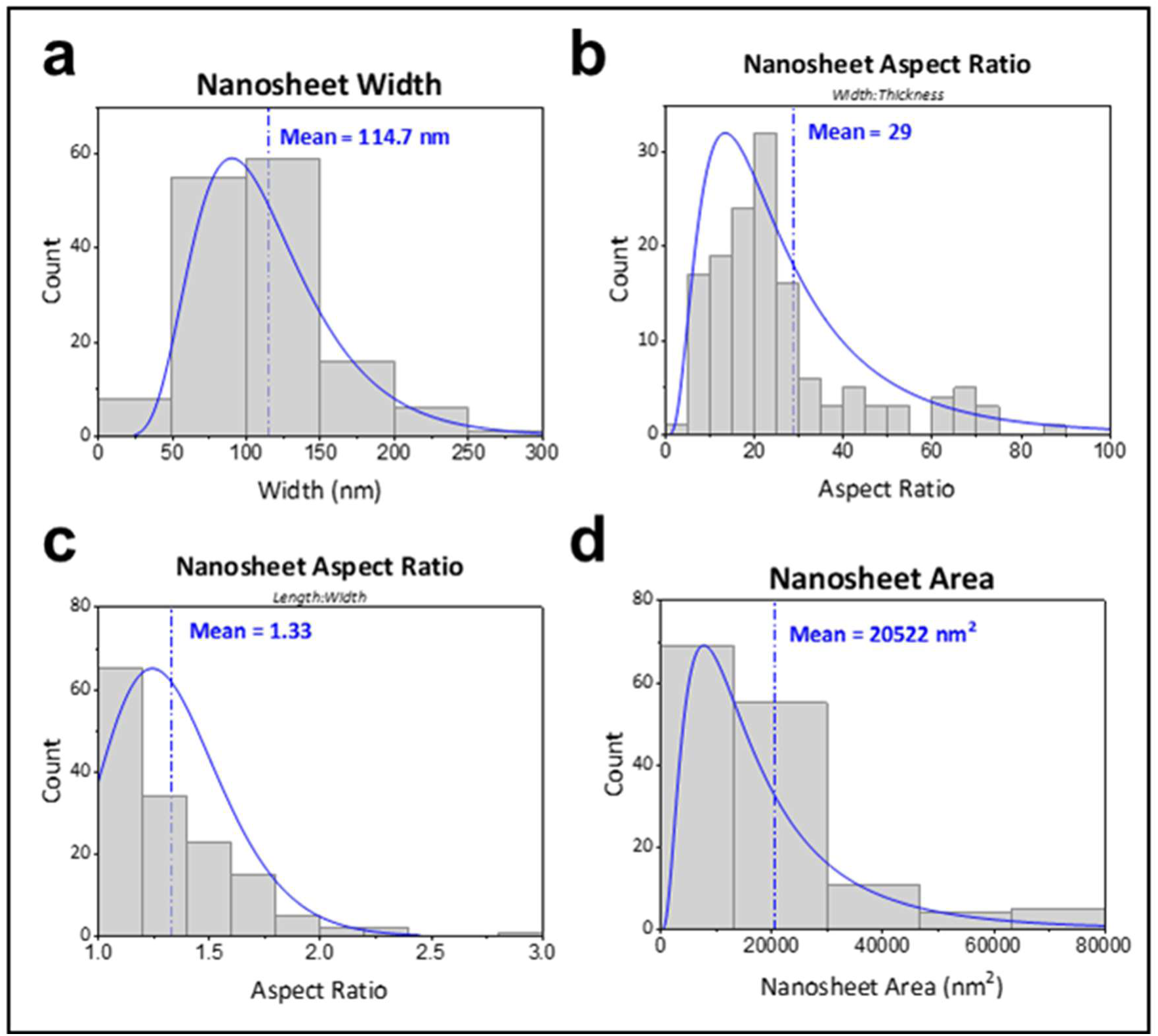
Further AFM characterisation of boron nanoplatelets: **a)** Nanosheet width **b)** Nanosheet aspect ratio (width:thickness) **c)** Nanosheet aspect ratio (length:width) **d)** Nanosheet area

**Fig. S3.**
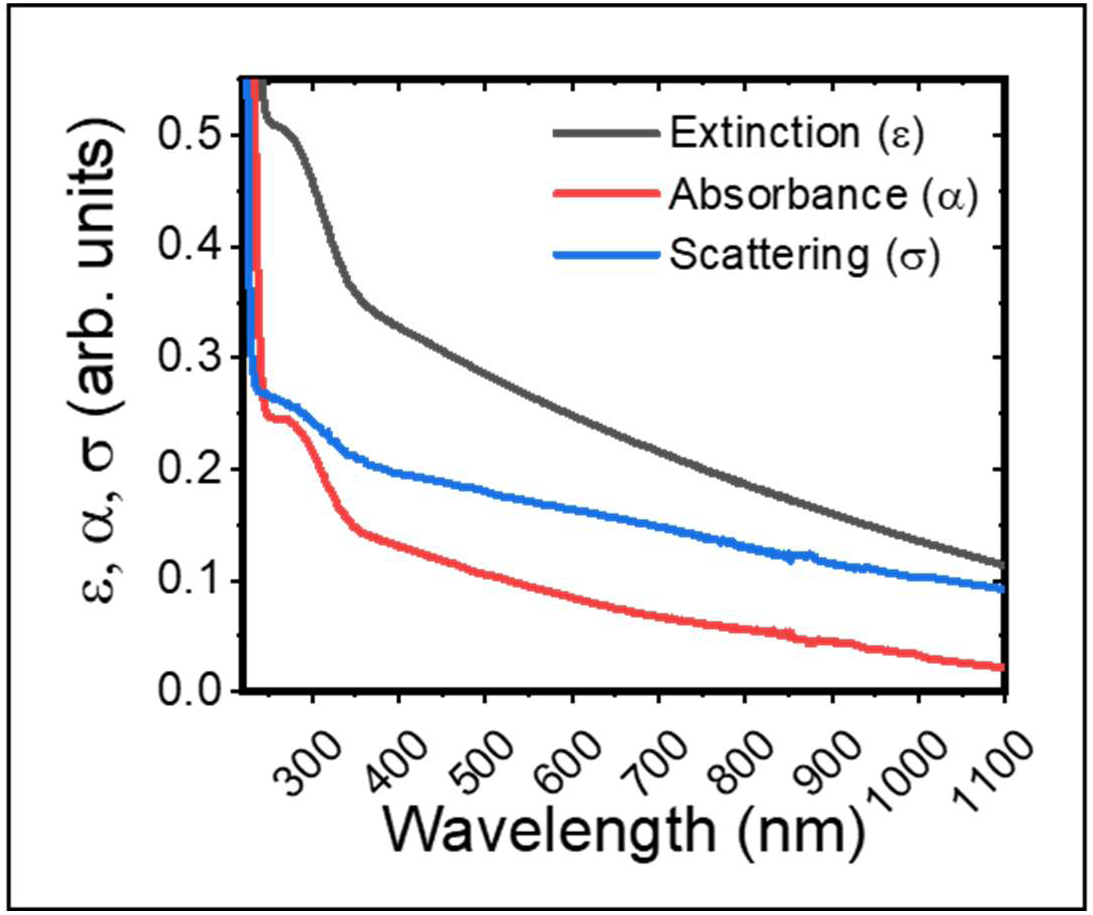
UV Visible light spectroscopy (UV Vis) of a boron nanosheet dispersion: Extinction (ε), scattering (σ) and absorption (α) spectra typical of a semiconducting nanomaterial

**Fig. S4.**
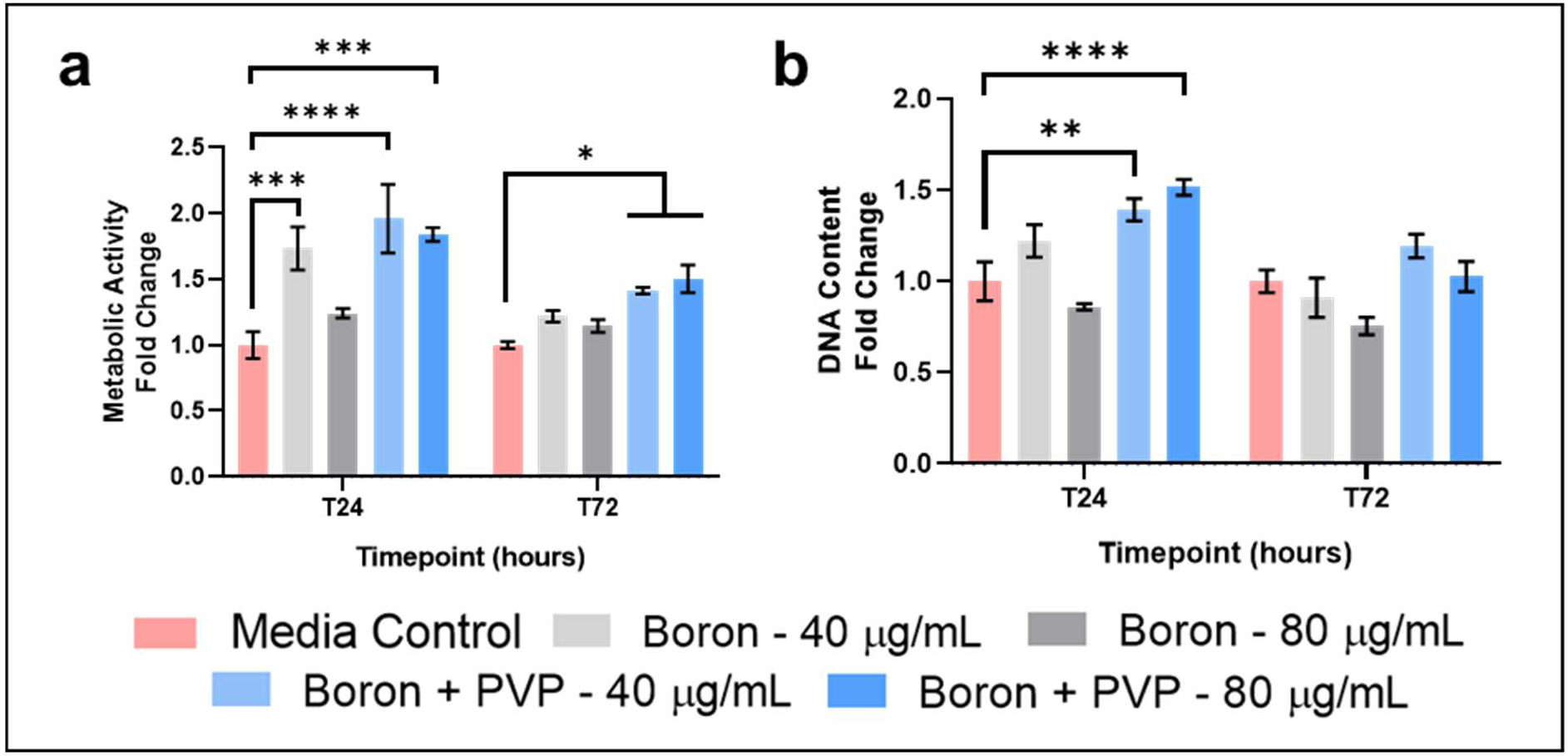
Biocompatibility of boron stabilization: a-b) It was necessary to determine if the nanosheets required any stabilization to remain dispersed and sufficiently hydrophilic in physiological conditions, as is the case with many nanomaterials of non-biological origin. To investigate this, mouse motor neuron cells (NSC-34 cell line) were grown in the presence of boron, with and without stabilization by PVP. This experiment demonstrated that PVP increased the biocompatibility of the boron nanosheets significantly, as seen by a significant increase in metabolic activity (a) and DNA content (b) for the PVP-stabilized material. Significances: *p < 0.05, **p < 0.01, ***p < 0.001, ****p < 0.0001

**Fig. S5.**
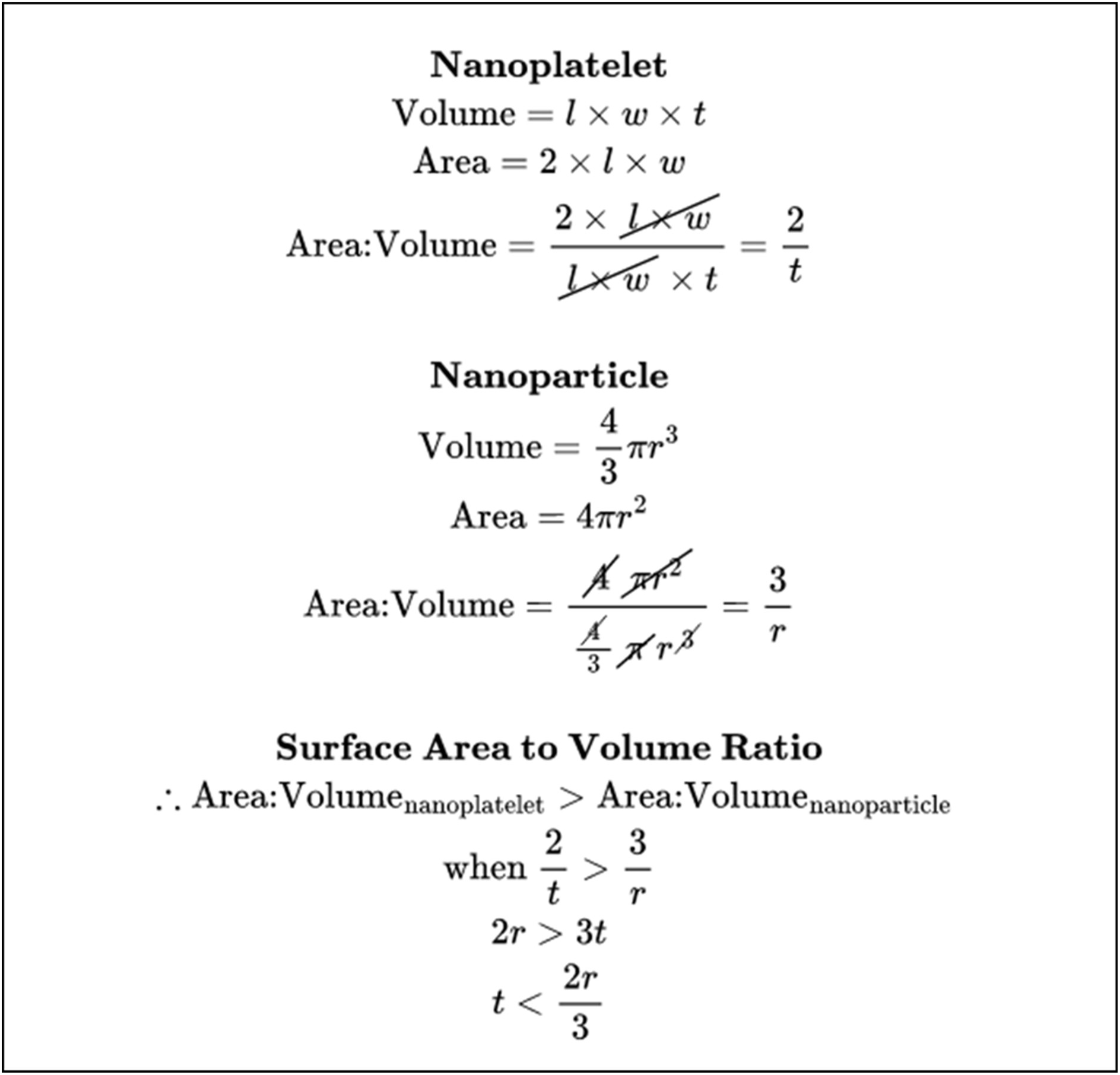
Derivation for the higher surface area:volume ratio of a nanoplatelet compared with a spherical nanoparticle: So long as the radius of the nanoparticle is >1.5 times the thickness of the nanoplatelet, the nanoplatelet will have a higher surface area:volume ratio

**Fig. S6.**
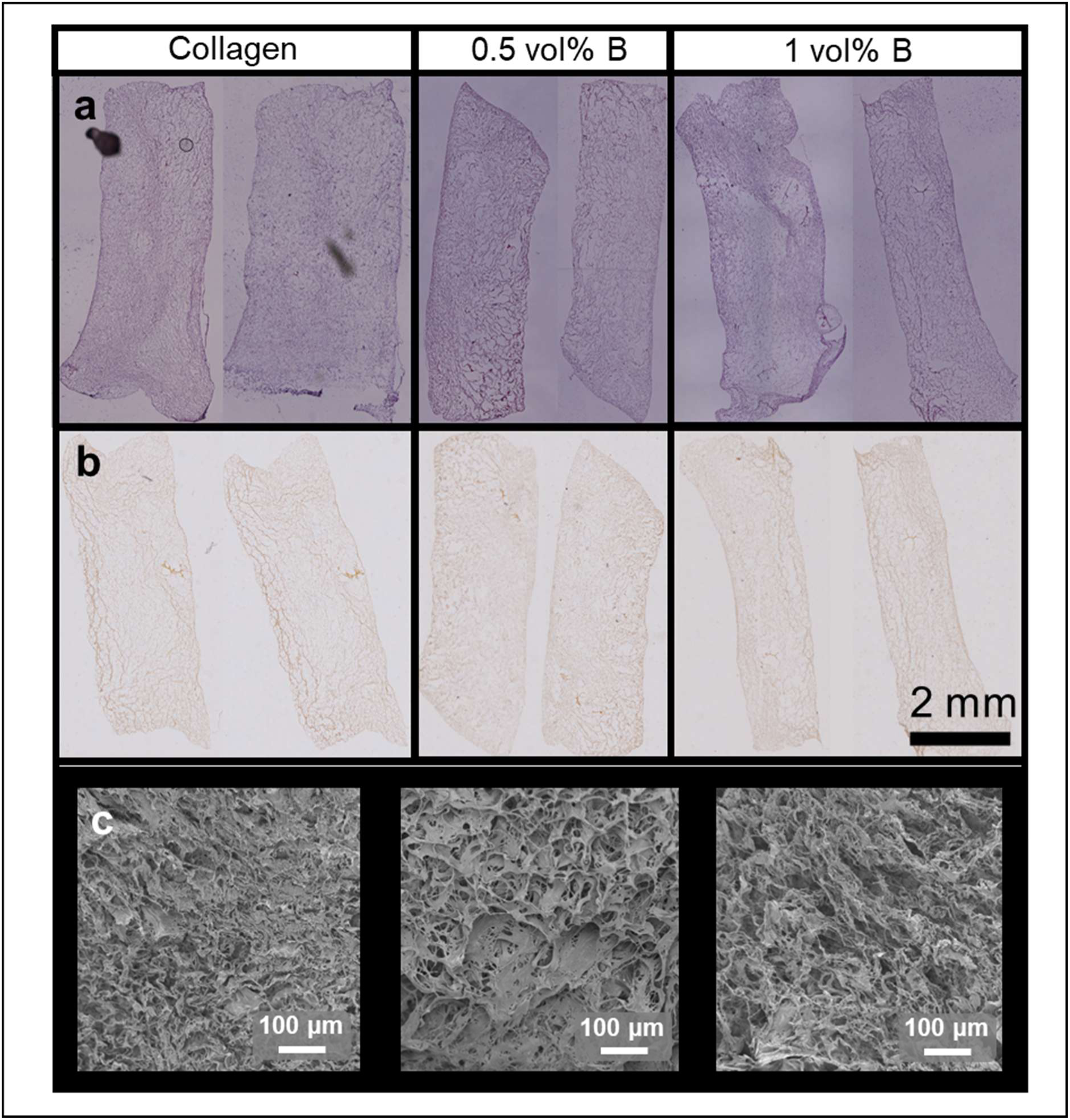
Histological staining of MC3T3s on BColl scaffolds: **a)** H&E histological staining of collagen-boron scaffolds with MC3T3s. **b)** Alizarin Red histological staining of collagen- boron scaffolds with MC3T3s. Scalebars for A & B - 2 mm **c)** SEM of MC3T3s in collagen- boron scaffolds. Scalebars 100 μm

**Fig. S7.**
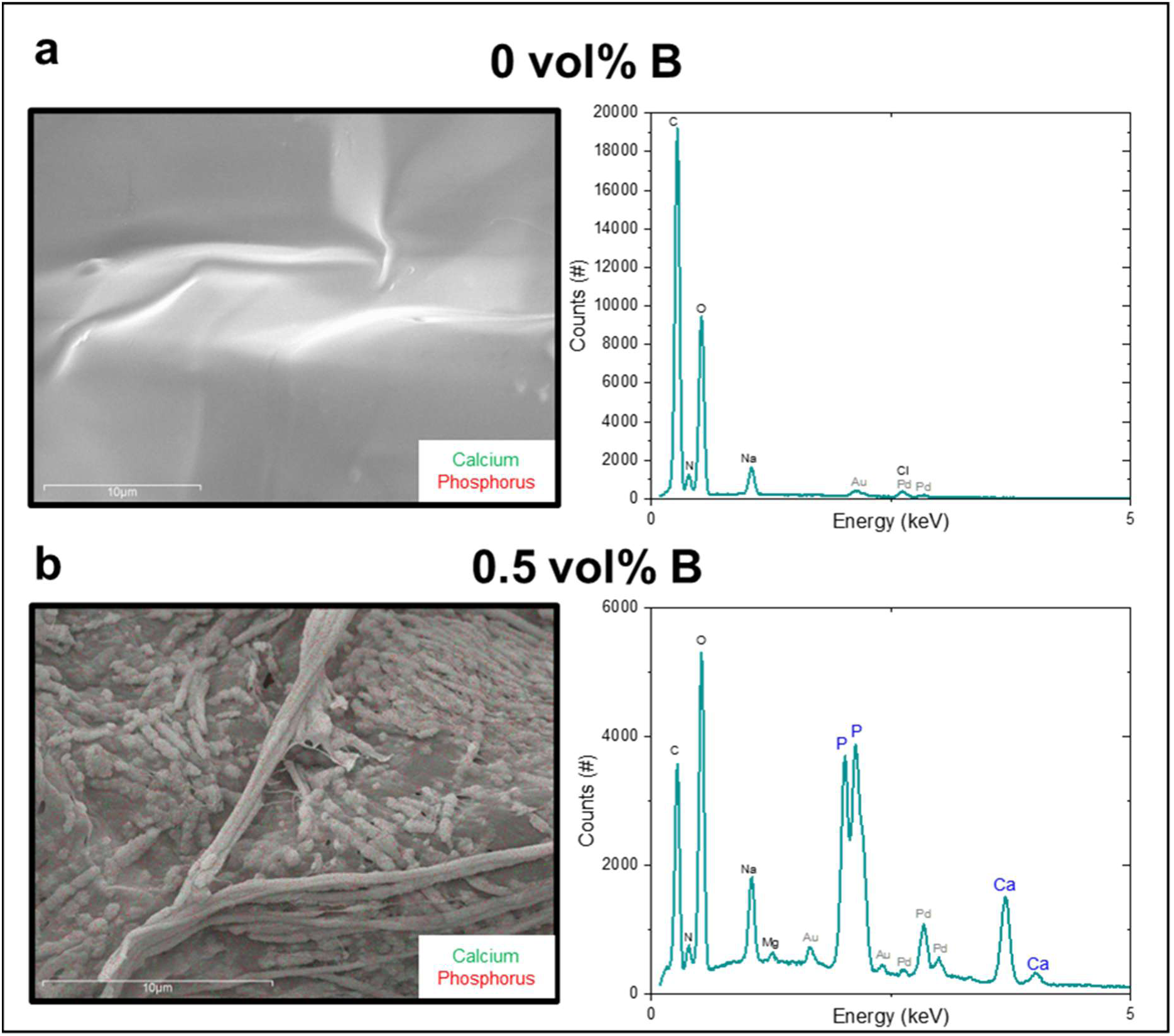
EDX of cell-laden BColl scaffolds: a-b) EDX analysis of collagen (a) and BColl (b) scaffolds, showing that the deposits on boron-containing scaffolds consist of calcium phosphate, the main component of bone mineral, corroborating enhanced bone mineralization in the presence of boron

**Fig. S8.**
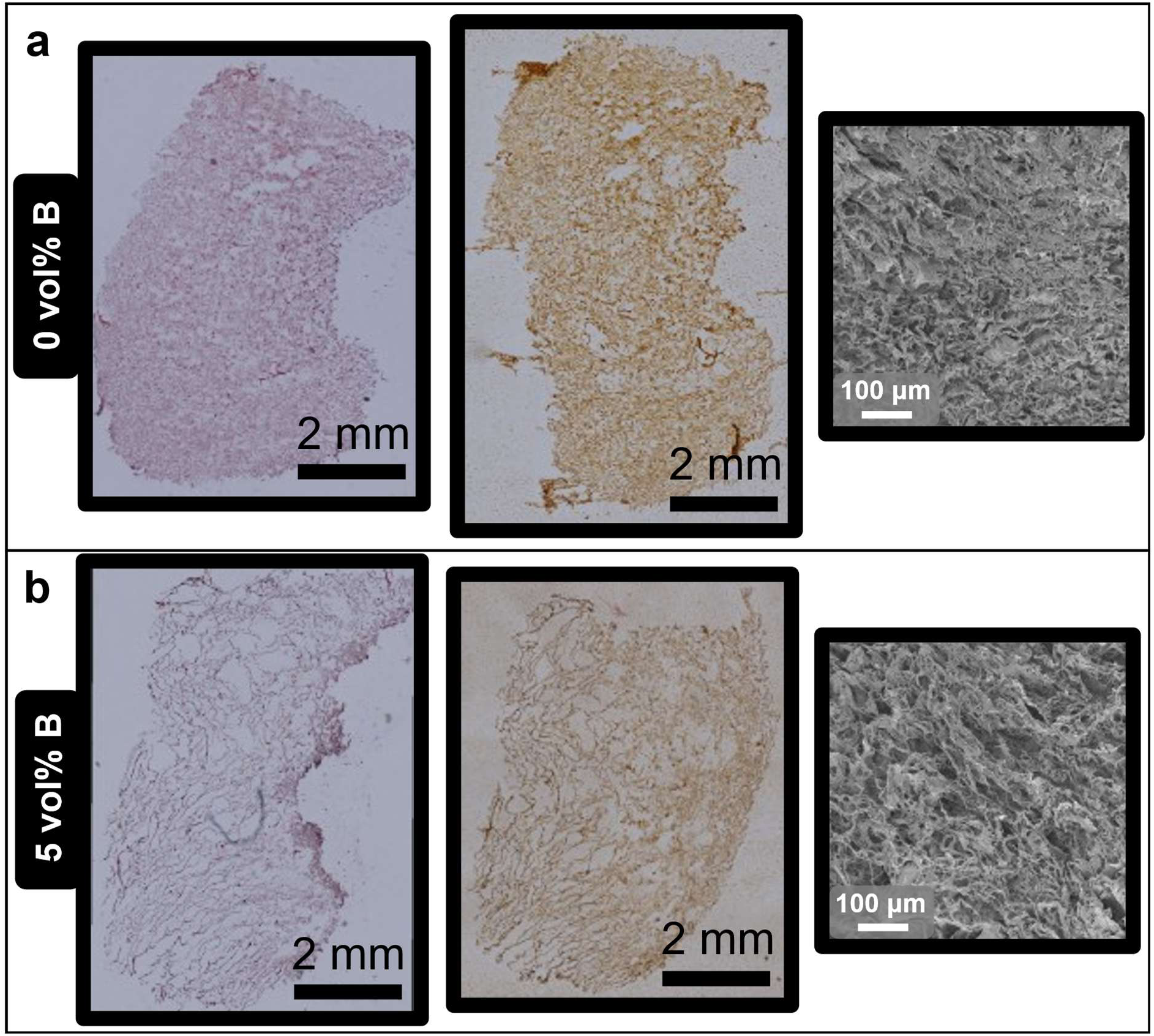
Histological analysis of rat mesenchymal stem cells on BColl scaffolds: a-b) To assess the extent and distribution of calcium deposition on the scaffolds, histological staining was carried out with H&E and Alizarin Red. Alizarin Red staining indicated robust mineralization of both collagen and BColl samples, throughout the sample. Scalebars 2 mm. SEM imaging was also carried out, to determine if there were any changes to the pore structure of the scaffolds following boron loading. Scalebars 100 μm

**Fig. S9.**
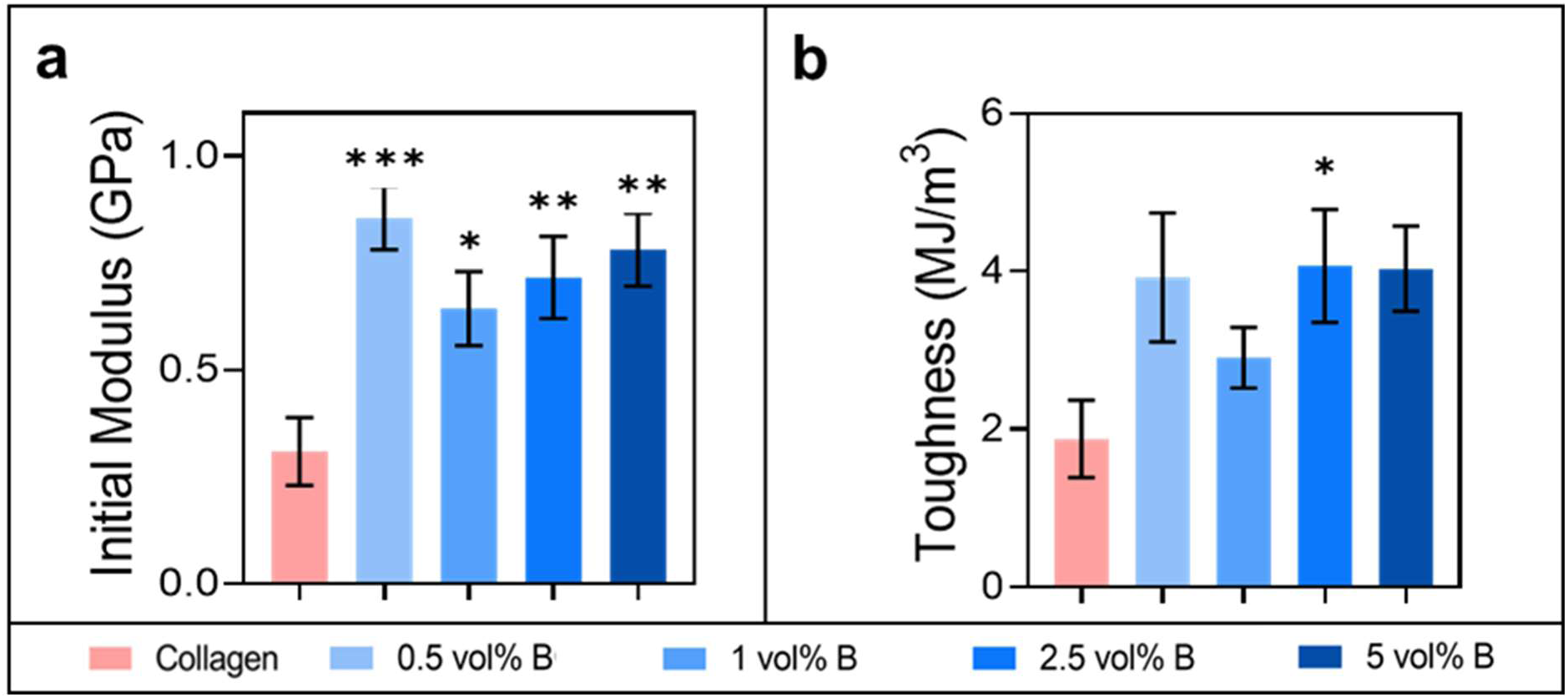
Further mechanical analysis of BColl films: **a)** Tensile modulus of BColl films in first 0.5% of strain, the regime most often experienced by cells, showing significant reinforcement in most BColl groups. **b)** Toughness measurements for BColl films, showing a trend towards increased toughness for all samples. Significances: *p < 0.05, **p < 0.01, ***p < 0.001, ****p < 0.0001

**Fig. S10.**
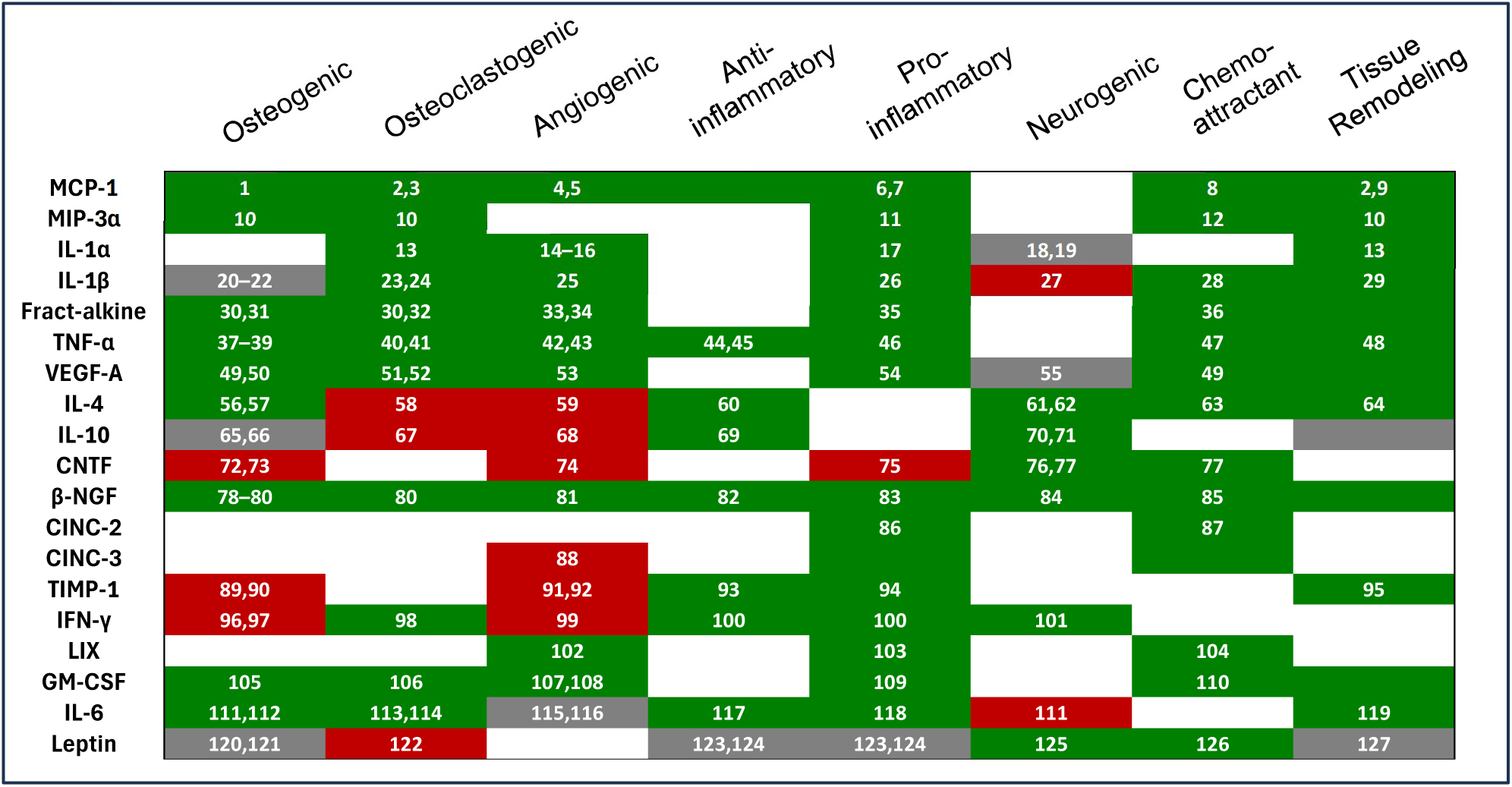
Investigation of paracrine release profile of rMSCs on BColl scaffolds: Matrix detailing the primary effects of each cytokine assessed in the ELISA data. Green corresponds to literature evidence of an effect, grey to contradictory or highly environment-dependent effect, red corresponds to evidence of a negative effect and white corresponds to no evidence for an effect. Numbers are citations

**Fig. S11.**
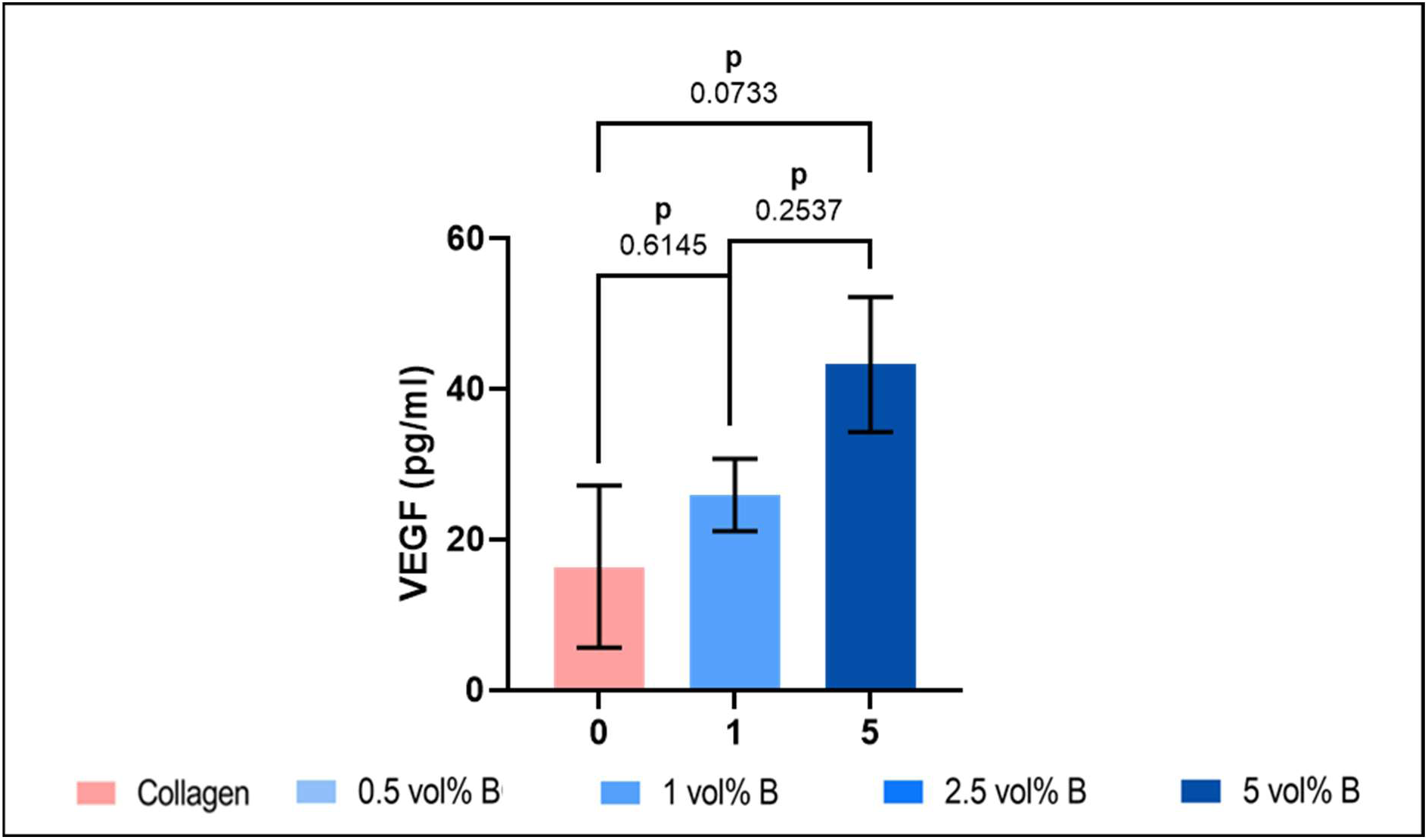
Assessment of angiogenic effect of boron: Analysis of VEGF release by human ELISA in rat MSCs after 3D scaffold culture for 28 days, showing trend towards increased VEGF release with boron addition

**Fig. S12.**
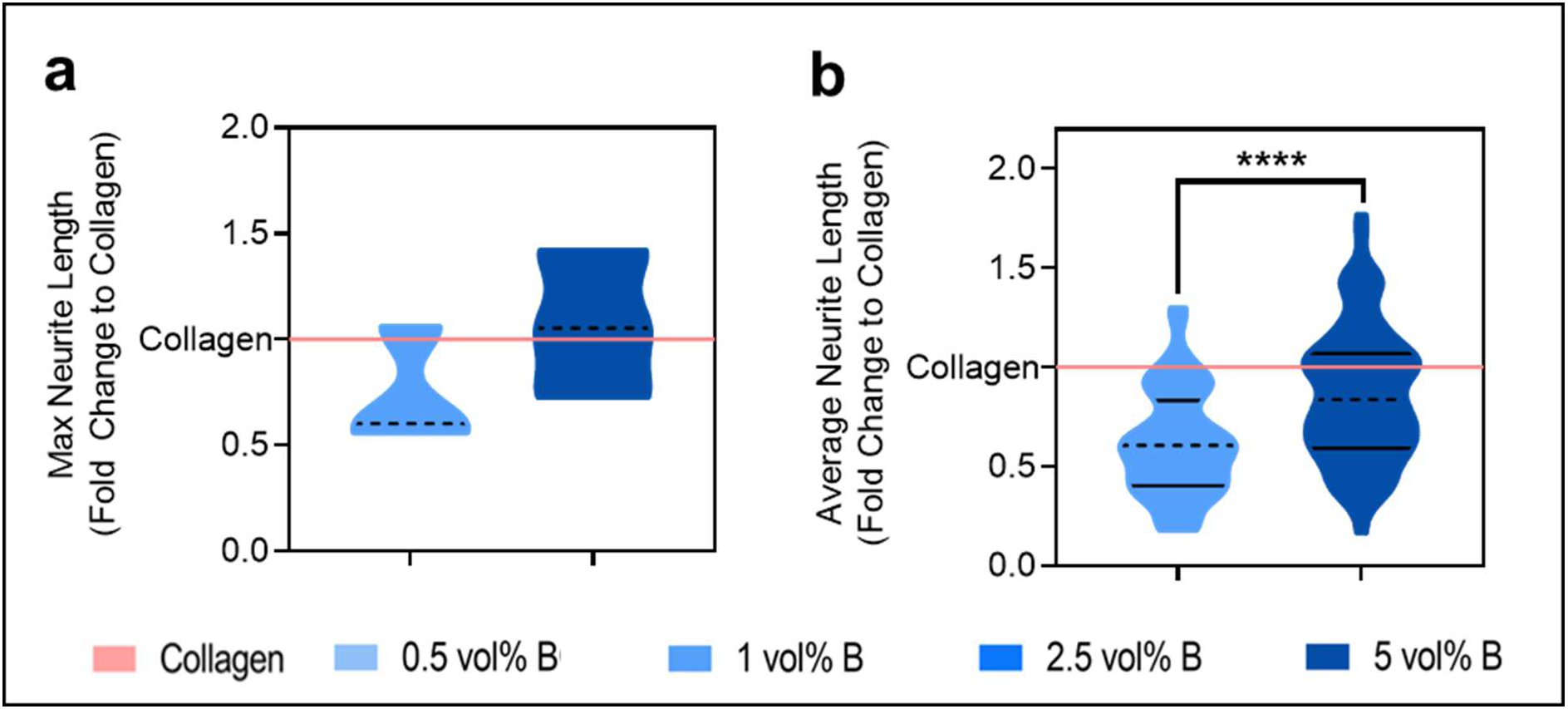
Assessment of neurogenic effect of boron: **a)** Max neurite length and **b)** average neurite length of DRGs grown on surface of BColl scaffolds for 14 days

